# Directing fratricide within T cell products using an anti-uPAR chimeric antigen receptor to drive the production of potent therapeutic cells

**DOI:** 10.1101/2025.10.24.684419

**Authors:** Lauren Sarko, David Givand, Claire Shepley, Brendan Rattin, Allen Attar, Rachel Taylor, Benjamin Kutler, Roshini M. Traynor, Anika Upadhyaya, Mackenzie Mnuk, Cavin Gehrke, Nat Murren, Tyler K. Ulland, Theresa Kotanchek, Krishanu Saha

**Affiliations:** Wisconsin Institute for Discovery, University of Wisconsin-Madison, Madison, Wisconsin, USA; Cellular and Molecular Pathology Program, University of Wisconsin-Madison, Madison Wisconsin, USA; Department of Biomedical Engineering, University of Wisconsin-Madison, Madison, Wisconsin, USA; Biomedical Research Model Services, University of Wisconsin-Madison, Madison, Wisconsin, USA; Department of Pathology and Laboratory Medicine, University of Wisconsin-Madison, Madison, Wisconsin, USA; Evolved Analytics LLC, Rancho Santa Fe, California, USA

**Keywords:** CAR, T cells, non-viral, CRISPR-Cas9, CD87, uPAR, scalable manufacturing, fratricide

## Abstract

Cell therapy manufacturing of primary T cells often results in heterogeneous cell populations within a final product, with many cells lacking desired of receptor expression or those that have exhausted or other dysfunctional phenotypes. Here, we design a novel cell-intrinsic strategy to genetically reprogram primary human T cells to autonomously detect and eliminate dysfunctional cells. This integrated detection and elimination process, known as directed fratricide, is programmed via nonviral CRISPR genome-editing to eliminate the T cell receptor (TCR) alpha chain (*TRAC* gene knockout) and integrate a chimeric antigen receptor (CAR) against the urokinase-type plasminogen activator receptor (uPAR), also known as CD87. Within these cell products, strong T cell stimulation or activation during manufacturing causes a small subset of cells to express uPAR, which subsequently triggers CAR-mediated killing by a separate subset of cells within the product. This fratricide induces proliferation in the desired cells and destroys undesired cells, a process that could be modeled computationally and controlled robustly via supplements to the culture media. The strategy enabled enrichment of anti-uPAR and anti-GD2 CAR T cell products up to ≥99% CAR+/TCR-, favoring a stem cell memory-like phenotype (CD45RA^high^/CD62L^high^). Understanding growth dynamics among T cell subsets and reprogramming them via CRISPR could accelerate the biomanufacturing of potent cell products without extensive selection methods.

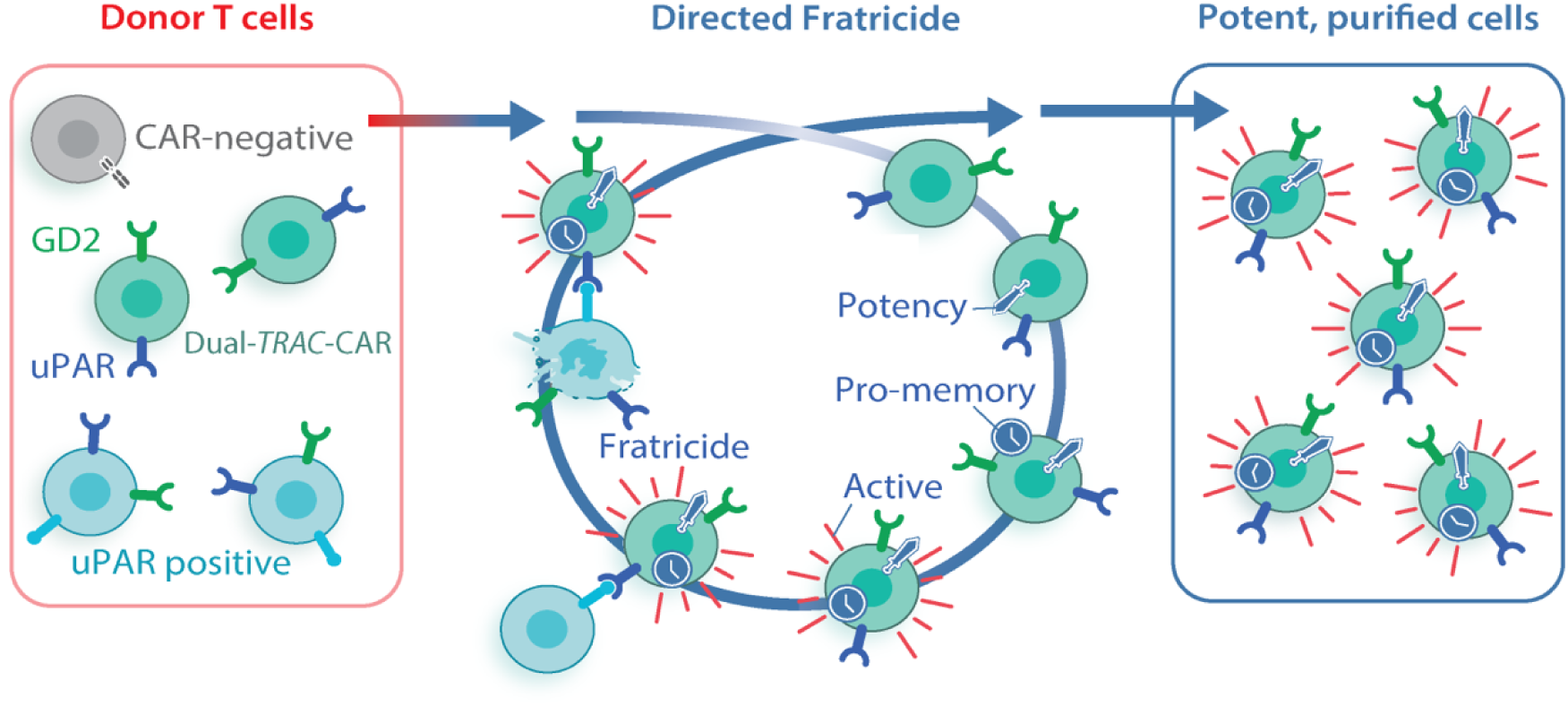

## Main Text

Genetically engineered T cells, like chimeric antigen receptor (CAR) T cells, can redirect T cell specificity and effector functions to attack a desired target cell. Achieving pure, potent cell products remains challenging due to incomplete transgene transfer and genome editing, especially for non-viral methods that are encumbered by inefficient delivery of large DNA templates, incomplete transgene knock-in, and donor-to-donor variability^1–5^. These constraints often result in heterogeneous transgene expression, requiring additional strategies to obtain therapeutically relevant cell populations^1,6–8^. Engineered T cells can be selectively enriched through co-expression of selectable markers such as drug resistance genes (e.g., dCK^9^, DHFR^10^), metabolic complementation systems^11^, inert surface markers (e.g., modified or truncated EGFR^12^, NGFR^13^, RQR8^14^), or fluorescent proteins, as well as by incorporating genetic modifications (cytokine autocrine loops^15^, anti-apoptotic signals^16,17^, immunosuppressive resistance^18^, base edited epitopes^19^) that confer a proliferative or survival advantage under defined culture conditions. In clinical manufacturing, surface marker-based FACS sorting and drug-selectable systems have been employed in some settings, with suicide genes (e.g., HSV-TK) or depletable markers enabling downstream safety control^20–24^.

Rather than relying on these strategies to enrich for potent engineered T cells, we sought to program cell-intrinsic signaling among T cells within the cell product to destroy dysfunctional cells autonomously. Establishing such a sensing and cytotoxicity selection strategy through genetic programming could simplify manufacturing and increase yields, as T cells are known to have intrinsic supernumerary capabilities *in vivo*^25^. In clinical CAR T manufacturing processes, a recent study found that a non-proliferating, senescent T-cell subpopulation in collections of autologous lymphocytes can have a negative impact on the manufacturing process^26^. Furthermore, CAR T cells have been directed previously to sense and eliminate senescent cells^27–30^. Notably, CAR T cells targeting the urokinase-type plasminogen activator receptor (uPAR, also CD87)^27,28,31,32^, can reverse senescence-associated pathologies, such as fibrosis and age-related metabolic disorders, within murine models. Within resting peripheral blood T cells, uPAR expression is undetectable and then rises upon T cell receptor (TCR) engagement with either anti-CD3/CD28/CD2 (TCR-dependent), PMA/Ionomycin stimulation in the presence of PP2 (TCR-independent), or stimulation with pro-inflammatory cytokines such as IL-2 or IL-7 ^27,33,34^. Upon TCR ligation *in vitro*, activation of the Src family kinases Lck and Fyn initiates downstream signaling cascades involving ZAP-70, LAT, PLCγ1, and PKCθ, culminating in the activation of transcription factors AP-1, NF-κB, and NFAT. This signaling cascade drives high-level uPAR expression, which has been suggested to facilitate T cell extravasation and migration^35,36^. Leveraging this activation-dependent expression pattern, we programmed a positive feedback loop to direct fratricide to selectively eliminate uPAR-positive T cells during *ex vivo* culture.

Our design uses CRISPR/Cas9-mediated genome editing^5,37–40^ to precisely integrate an anti-uPAR CAR to eliminate undesired T cells in the cell product that arise during manufacturing. Directed fratricide in these cell products was modeled through symbolic regression which predicted culture conditions that result in cell products highly enriched for stem cell memory-like phenotypes and transgenes (e.g., ≥99% CAR+). The facile biomanufacturing of CAR T cell products with these attributes could enable a new class of immune cell therapies that encode selective purification mechanisms within cells through precise genome editing and receptor design.

## Results

### Enabling a directed fratricide via anti-uPAR CAR and CRISPR

We utilized homology-directed repair (HDR) after CRISPR/Cas9 cleavage of DNA to integrate an anti-uPAR CAR precisely at the start of the first coding exon of the *TRAC* locus (Figure 1A). Physiological expression of CARs integrated at this site was previously characterized, specifically with an anti-CD19 CAR^40^ and anti-GD2 CARs^5,37,38^. Off-target analysis for this single guide RNA (sgRNA) has been well characterized, with minimal off-target editing^37,41^. For the HDR template, we synthesized a second-generation uPAR-CAR sequence with an mCherry fluorescent protein and inserted it into a Nanoplasmid backbone^4,27,28,42^. The mCherry is separated from the CAR through a self-cleaving 2A peptide, facilitating the imaging and detection of CAR-expressing cells. The Nanoplasmid backbone reduces the length of the plasmid backbone, as delivery of linear double-stranded DNA (dsDNA) can generate toxicity when delivered into T cells^43^. Our uPAR-CAR template has a single chain variable region (scFv) against the human uPAR protein antigen and human costimulatory domains CD28 and CD3σ (uPAR-*TRAC*-CAR).

**Figure 1.**
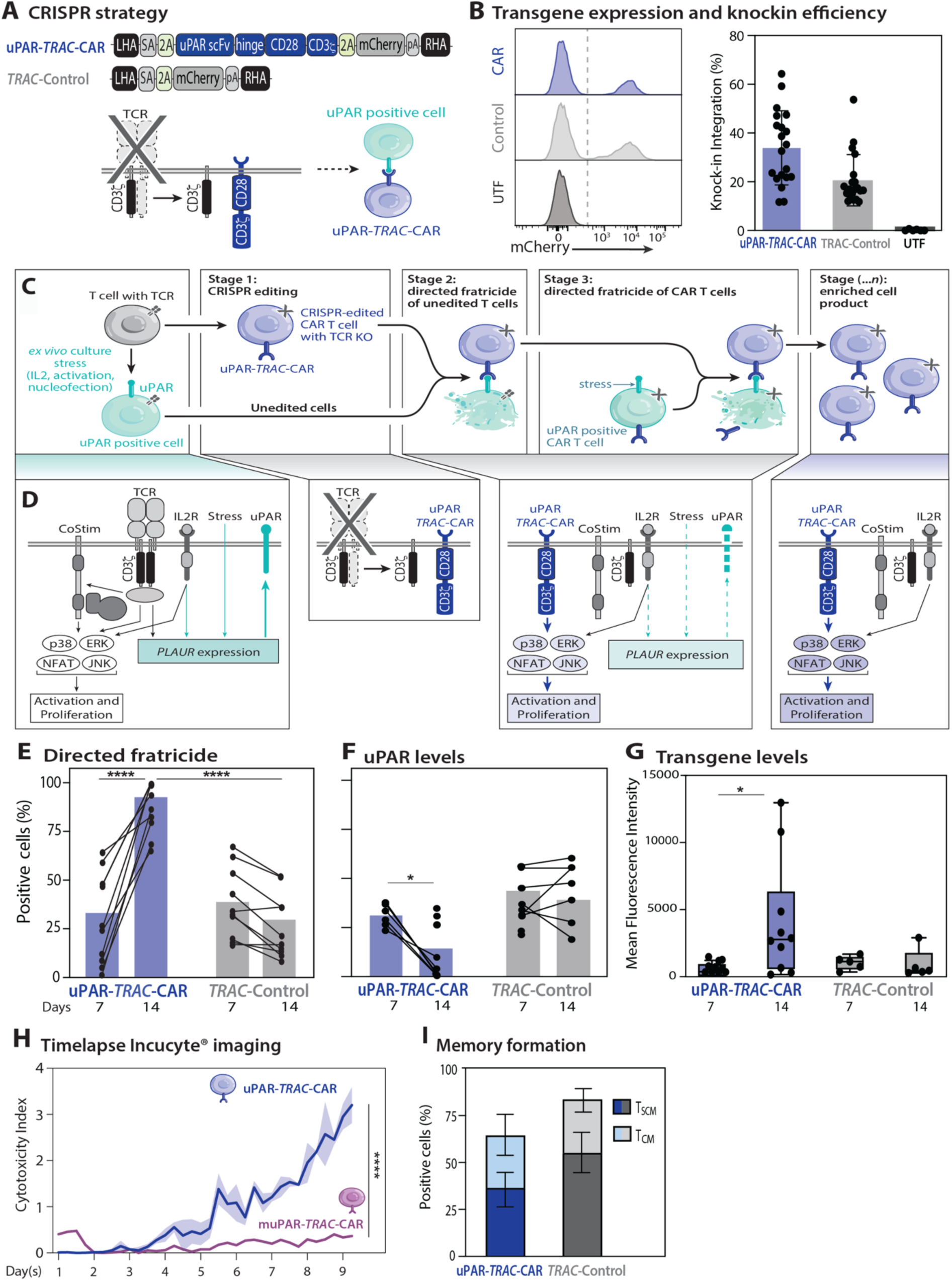
Design and evaluation of directed fratricide in primary human T cells. (A) Schematic of uPAR-*TRAC*-CAR and *TRAC*-Control construct targeting the first encoding exon of the human *TRAC* gene consisting of a SA: splice acceptor (grey), T2A: self-cleaving peptide (green), uPAR scFv: single chain variable fragment targeting human uPAR (blue), hinge domain, human costimulatory domains: CD28 and CD3ζ (blue), 2A: self-cleaving peptide (green), mCherry: fluorescent marker (grey), pA: rabbit ß-globin polyA terminator (grey). (B) Histograms of %mCherry expression and summary of flow cytometry knock-in and knock-out of the manufactured cell products on day 7 post-isolation. uPAR-*TRAC*-CAR (blue, n=20), *TRAC*-Control (grey, n=20), Untransfected (UTF) (black, n=6) over 6 donors. (C-D) Cell signaling crosstalk schematic within uPAR-*TRAC*-CAR T cell culture enables directed fratricide. Early stages of directed fratricide differentially activate *PLAUR* and initiate a cascade of uPAR-CAR activity against uPAR positive T cells in the culture vessel. (E-F) Summary of flow cytometry for (E) %CAR, and (F) uPAR in T cell cultures before and after a seven day directed fratricide period: day 7 and 14 post-isolation in *ex vivo* T cell culture. uPAR-*TRAC*-CAR (blue, n=10), *TRAC*-Control (grey n=10) across three donors. (G) Mean Fluorescence intensity (MFI) of %CAR and %mCherry of uPAR-*TRAC*-CAR and *TRAC*-Control T cells during 7 day directed fratricide period. uPAR-*TRAC*-CAR (blue, n=10), *TRAC*-Control (grey, n=10) across three donors. (H) IncuCyte® Live-Cell Analysis of total red object integrated intensity (RCU) cytotoxicity index and proliferation of uPAR-*TRAC*-CAR (blue) and muPAR-*TRAC*-CAR (purple) showing a significant increase in fluorescence intensity during directed fratricide. (I) Summary of flow cytometry immunophenotyping showing percent positive of T cell differentiation states: T_SCM_ CD45RA+/CD62L+ and T_CM_ CD45RO+/CD62L+ for both uPAR-*TRAC*-CAR (blue; n=5) and *TRAC*-Control (grey;n=5) on day 14 post-isolation across three donors. Significance was determined by (E, F, H) two-way ANOVA, (G) ordinary one-way ANOVA, (I) mixed ANOVA; *p≤0.05; ***p≤0.001;****p≤0.0001. ANOVA, analysis of variance.

Primary human T cells from healthy donors were electroporated with the purified HDR templates and SpCas9 ribonucleoproteins (RNPs) targeting the human *TRAC* locus. Two different HDR templates were investigated: the uPAR-*TRAC*-CAR template and a no CAR control template (*TRAC*-Control), which contains only the mCherry reporter gene in place of the uPAR-*TRAC*-CAR sequence. During the first two weeks of manufacturing, the cells were assayed for cell count and viability (Figure S1A-B). A noticeable decrease in viability and cell count was observed post-nucleofection, likely due to the stress from electroporation induced membrane disruption along with the large, multiple kilobase size of the dsDNA templates. Despite this transient decrease, we achieved consistently high gene editing with the dsDNA templates across eleven different donors and demonstrated up to 65% knock-in efficiency, with an average of 23% for uPAR-CAR+ (assayed via mCherry reporter expression) and >90% total TCR- cells, as measured by flow cytometry on day 7 of manufacturing (Figure 1B). On-target genomic integration of uPAR-*TRAC*-CAR was confirmed via “in-out” PCR amplification assay on genomic DNA extracted from donor-matched samples with primers specific to the *TRAC* locus outside the left homology arm and inside the CAR transgene (Figure S1C). On day 11 of manufacturing, we evaluated the cytotoxicity and function of uPAR-*TRAC*-CAR T cells against both irradiation-induced senescent uPAR+ cells and a uPAR- overexpressing HEK-293 (uPAR-293) cell line. We observed robust potency against these uPAR- expressing target cells within 48 hours of co-culture (Figure S2).

Within our cell products, directed fratricide during uPAR-*TRAC*-CAR T cell manufacturing was evaluated on day 7 and day 14 post-isolation. Both the frequency of uPAR-*TRAC*-CAR T cells and the mean levels of CAR on the cell surface increased significantly over time, indicating growth of CAR-expressing cells and upregulation of CAR expression in many cells. No comparable increase was observed in *TRAC*-Control T cells (Figure 1E, 1G). To determine if this increase in CAR expression was due to target antigen exposure, we measured uPAR expression via flow cytometry. uPAR was detectable in a subset of T cells (20-60%) in both cultures on day 7, but on day 14, there was a significant decrease in uPAR-positive cell subpopulations in uPAR-*TRAC*-CAR T cell products (Figure 1F). There was no decrease of this subset in *TRAC*-Control T cells, indicating that the reduction of uPAR-positive cells was CAR-dependent (Figure 1C-D). To confirm that CAR signaling was controlling directed fratricide, we utilized the tyrosine kinase inhibitor, dasatinib, to block lymphocyte-specific protein tyrosine kinase (Lck) and prevent CD28-based signaling downstream of CAR activation^44–46^. We treated uPAR-*TRAC*-CAR T cells with 100 nM dasatinib over several time courses. The duration of dasatinib-based inhibition of CAR signaling (Figure S3A) correlated with the percentage of uPAR-*TRAC*-CAR in the final product, with more undesired CAR- negative cells persisting in the culture when CAR signaling was inhibited (Figure S3B).

To further assess whether the directed fratricide is due to the antigen binding from our CAR, we exchanged the scFv in the uPAR-CAR with a scFv specific for murine uPAR (muPAR), because the epitope on uPAR for the human scFv is not conserved in mice^47,48^. We generated a Nanoplasmid HDR template with a modified scFv to enable binding to the murine uPAR antigen (Figure S4A)^13^. As above, primary human T cells from healthy donors were electroporated with the purified HDR templates and Cas9 RNPs targeting the human *TRAC* locus. The process for muPAR-*TRAC*-CAR production was highly reproducible across six donors and yielded potent muPAR-*TRAC*-CAR T cells against murine uPAR-positive target cells (Figure S4B-E). After confirming the functionality of muPAR-*TRAC*-CAR T cells, we manufactured donor-matched uPAR-*TRAC*-CAR and muPAR-*TRAC*-CAR products and cultured them in parallel imaging each condition independently throughout the directed fratricide period post-electroporation. We observed a striking increase in the desired transgene-positive T cells throughout the observation period only in the uPAR-*TRAC*-CAR T cell products (Figure 1H). In contrast, the muPAR-*TRAC*-CAR T cells maintained the same fraction of desired cells, without any elimination of undesired cells. This increase in the desired uPAR-CAR-containing T cell fraction indicated that directed fratricide in the uPAR-*TRAC*- CAR T cell products is a result of the CAR binding to human uPAR antigen.

Tumorigenic potential and off-target activation of our *TRAC*-based uPAR-CAR knock-in strategy were assessed *in vivo*. Current regulatory guidance for cell therapy products^49^ with gene-edited cells or CARs recommends evaluating safety through the characterization of cytokine-independent growth and off-target activation. We assessed whether antigen-stimulation *in vivo* within mice would promote tumor formation, cytokine release syndrome, or other toxicity by administering muPAR-*TRAC*-CAR cell products into mice. We chose to infuse into the brains of adult immunodeficient mice via intracerebral infusion to mimic the potential serious off-target activation of a uPAR-*TRAC*-CAR cell product in an adult patient. After dosing with up to 500,000 CAR-positive T cells, none of the mice had significant changes in weight or temperature (Figure S4G-H), and event-free survival was 100% (Figure S4F).

### Phenotypes of uPAR-*TRAC*-CAR cell products

Chronic activation during manufacturing can induce tonic signaling^50^, which may lead to excessive differentiation and reduced efficacy upon antigen engagement. In contrast, *TRAC*-CAR T cells retain a memory-like phenotype following antigen exposure in co-culture^37,38,40^. Because of the significant antigen exposure during *ex vivo* manufacturing, this memory-like phenotype could differentiate into undesired exhausted cells after directed fratricide. Therefore, we tracked cell fates using immunophenotyping via flow cytometry for markers of T cell memory and exhaustion at the end of manufacturing on day 14 (Figure 1I; S1D). First, we assessed the expression of T cell naïve/memory markers, CD45RO, CD45RA, and CD62L. At day 14 despite uPAR antigen exposure, uPAR-*TRAC*-CAR T cells still consist of a predominantly (37% T_SCM_) stem cell memory phenotype and (30% T_CM_) central memory phenotype (Figure 1I). This phenotype post-antigen exposure suggests that our uPAR-*TRAC*-CAR T cell products still have high potency *in vivo* based upon prior studies of memory cells in patients, which have a high capacity to self-renew and persist long-term in patients^51^. As expected, a subset of T cells differentiated with expression of CD45RA, CD45RO, and CD62L at the end of manufacturing, aligning with our previous findings that culturing *TRAC*-CAR T cells with Immunocult XF medium and IL-2 can generate high expression of CD45RA and CD62L^14^. Second, we also profiled three key immune checkpoint molecules, PD-1, TIM-3, and LAG-3. The co-expression of these three inhibitory receptors (IRs) can indicate a state of exhaustion, making T cells less functional and unable to eliminate the antigen of interest. Expression of all three exhaustion markers in uPAR-*TRAC*-CAR and *TRAC*-Control T cells were comparable (Figure S1D). Moderate TIM- 3 expression is expected due to the consistent stimulation of the CAR due to chronic antigen exposure and strong cytokine signaling via IL-2 media supplementation^52,53^. Despite our uPAR-*TRAC*-CAR T cells displaying an overall CD45RA^high^/CD62L^high^ population and low IR co-expression, we observed a small subset of uPAR+ cells at the end of manufacturing, indicating cell dysfunction due to chronic antigen signaling (Figure 1F). Further elimination of this cell subset was the focus of our following experiments.

### Modeling directed fratricide to increase memory phenotypes

To determine what variables impact uPAR expression during manufacturing, we used DataModeler, an augmented intelligence platform that utilizes multi-objective symbolic regression to generate model ensembles of the highest performing variable combinations. Culture conditions, cell growth, cell viability, and cell type subsets were integrated into a data table with 1273 samples across many manufacturing runs. Subsets of the data table were evaluated and analyzed to determine what factors affected the desired cell fraction (percentage of CAR-positive cells, %CAR) in culture over time in the uPAR-*TRAC*-CAR products. Pairwise linear correlation plots were created to determine which variables within our data table could influence %CAR as well as uPAR expression. Through variable correlation analysis, we determined that there was a positive correlation between %CAR and units of IL-2 added to the culture media (Figure 2A). uPAR expression also correlated negatively with memory marker, CD45RA, and positively with exhaustion markers PD-1 and LAG3 (Figure 2A; S5A-B).

**Figure 2.**
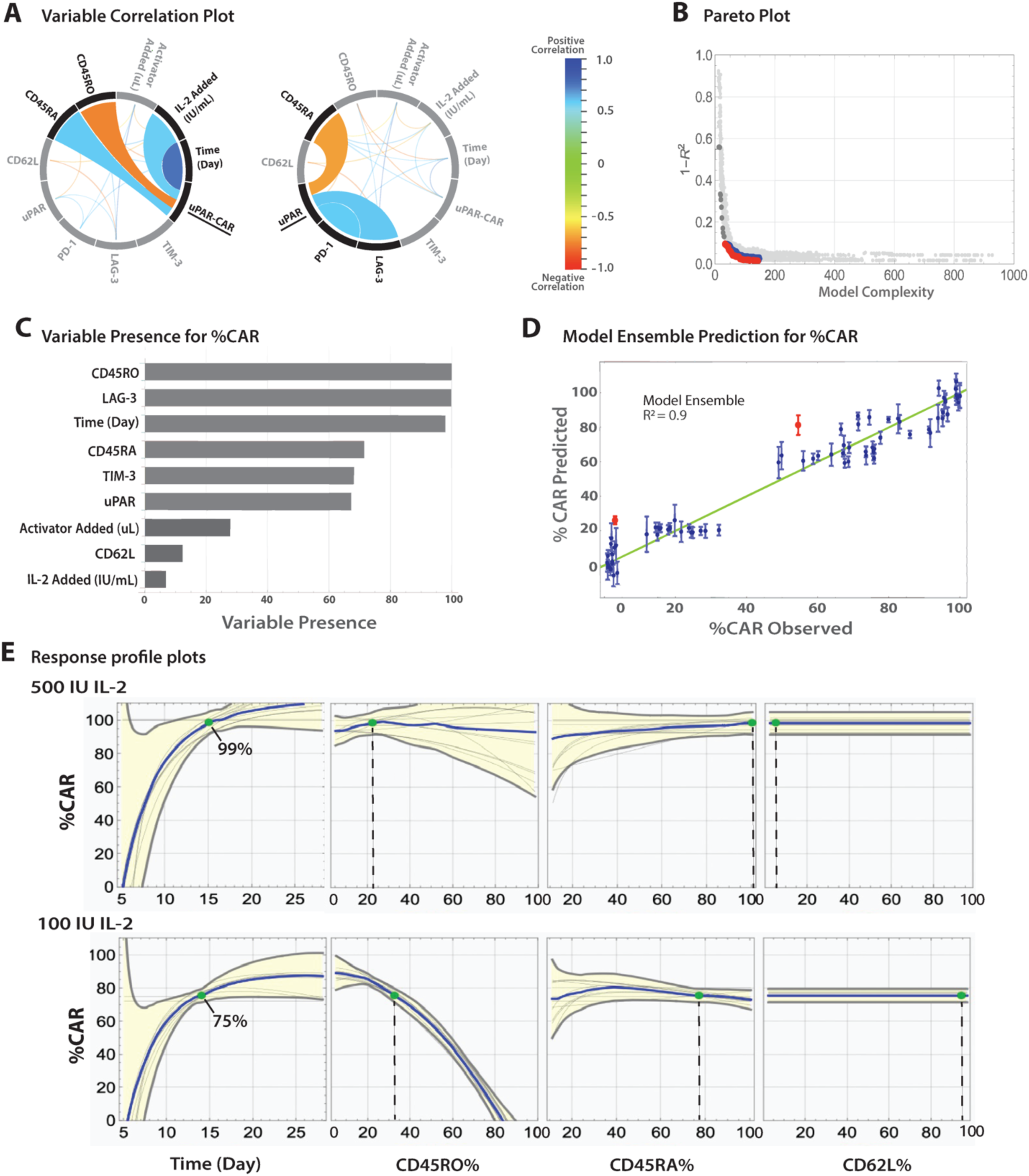
Nonlinear regression analysis shows a relationship between culture conditions, %CAR, and T cell phenotype. (A) Variable correlation plots for uPAR-CAR and uPAR levels among samples collected during manufacturing. (B) Pareto plot identifying the top 896 models selected at the knee of the Pareto Front. Red models distinguish the most accurate models (R^2^>0.9) with the lowest degree of model complexity on the Pareto Front, while blue dots represent other models within the quality box (R^2^>0.9, complexity≤150). (C) Variable presence chart showing the most common variables present in the models at the Pareto front. (D) Model ensemble prediction plot demonstrating high predictive power of the model ensemble for the observed data. The error bars identify the 2σ spread of the ensemble predictions at each data point. (E) Response profile plots of the model ensemble for %CAR for time, IL-2, and phenotype at 500 IU IL-2 and 100 IU IL-2/mL culture supplementation with 0 μL activator. The gray lines represent individual models, while the blue line represents the median of the predictions of the ensemble. The yellow shaded area marks the 2σ spread of the ensemble prediction.

Models were then generated using the following variable inputs: time in culture, IL-2 added, activator added, phenotype markers (CD45RO, CD45RA, CD62L), and exhaustion markers (PD-1, LAG3, TIM-3). Through several rounds of modeling, we found that models with low complexity – incorporating only 5, 6, or 7 dimensions - could predict the %CAR over time with high accuracy (*R^2^*>0.9, Figure 2B). The variables and variable combinations present in the high-accuracy, low-complexity models were measured and ranked (Figure 2C). The most common variables in the selected models were used to further investigate the factors that impact uPAR and %CAR expression.

Using six selected variables, we generated model ensembles to create predictive models of uPAR-*TRAC*- CAR (Figure 2D; S5C). Our modeling identified that cell products exposed to higher levels of IL-2 and activator were more likely to co-express the IR markers (Figure 2E; S5D-F). The model predicted that reducing overall uPAR levels by reducing IL-2 addition to the culture media would increase %CAR and retain memory markers (Figure 2E).

Guided by the modeling results, we performed new experiments with primary T cells under different culture conditions. First, we activated the cells with 200 IU IL-2 and the standard 25 µL/mL CD28/CD3/CD2 activator at a density of 2 million cells/mL. After electroporation to introduce either uPAR-*TRAC*-CAR or *TRAC*-Control transgenes, the T cells were maintained in culture with either 100 or 500 IU IL-2/mL. On day 14, uPAR-*TRAC*-CAR T cells cultured with 100 IU IL-2 had a significant decrease in uPAR expression compared to cells cultured in 500 IU IL-2 (Figure 3A). To help balance uPAR expression and provide enough antigen for directed fratricide in the culture, we manufactured our uPAR-*TRAC*-CAR and *TRAC*- Control T cells as described above with 100 IU IL-2 and either 0 µL/mL, 25 µL/mL, or 50 µL/mL CD3/CD28/CD2 activator. While %CAR for uPAR-*TRAC*-CAR T cells was not affected by these different activator conditions (Figure S5G), we observed a significant increase in %CAR in uPAR-*TRAC*-CAR T cells when both activator and IL-2 levels were lowered together, whereas the %mCherry transgene expression in *TRAC*-Control T cells decreased over the same time period (Figure 3B-C). Importantly, a high percentage of pro-memory phenotypes (>80% with T_SCM_ or T_CM_ markers) was preserved with reduced IL-2 and activator culture conditions, supporting our model ensemble plot predictions (Figure 3D).

**Figure 3.**
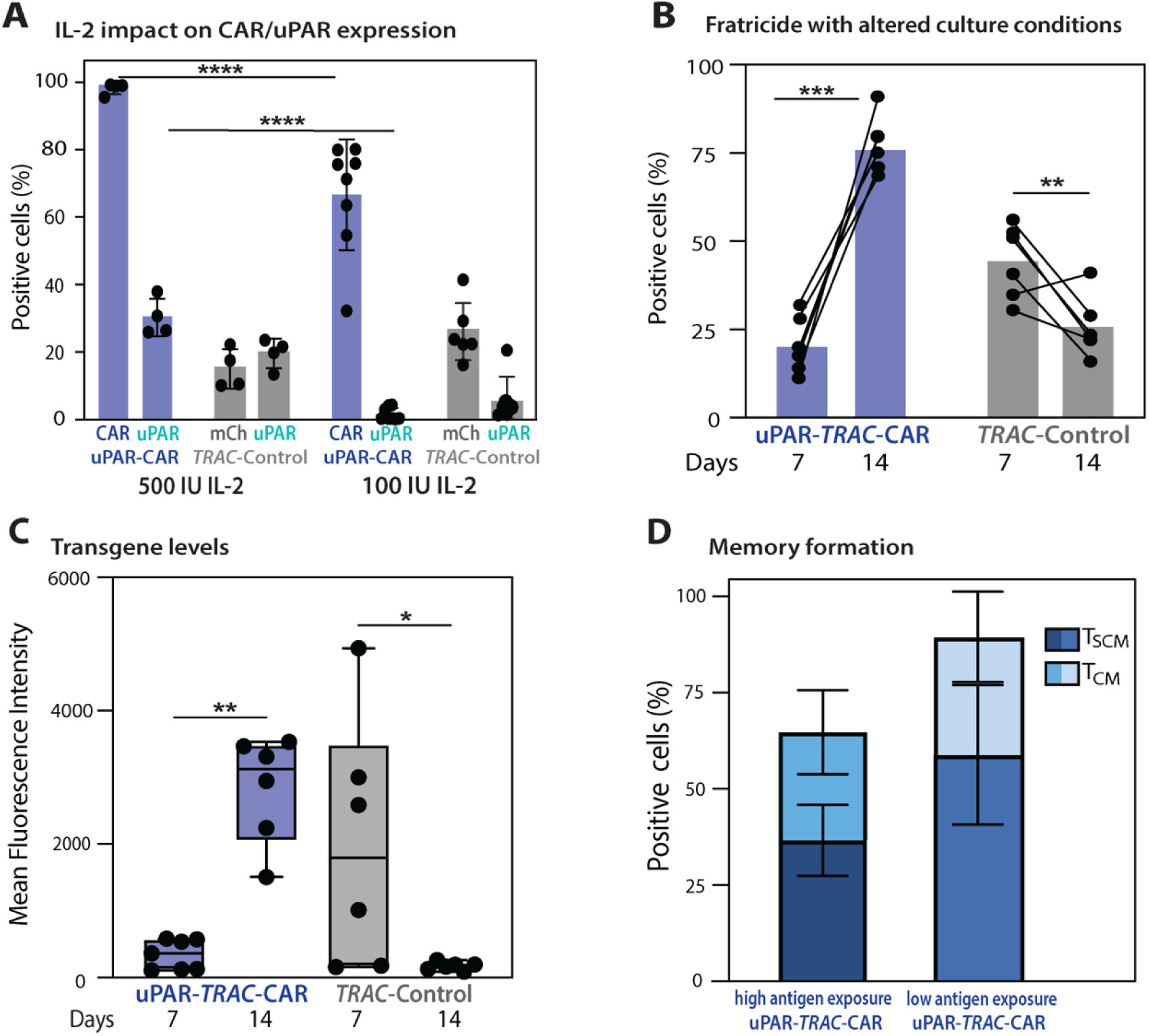
Predicted culture conditions preserve memory-like phenotype after directed fratricide of uPAR-*TRAC*-CAR T cells. (A) Flow cytometry of %CAR, %mCherry, and %uPAR expression for uPAR-*TRAC*-CAR (blue) and *TRAC*-Control (grey) on day 14 of CAR T cell manufacturing after being cultured with either 500 or 100 IU IL-2/mL. uPAR-*TRAC*-CAR showed a significant decrease in both CAR and uPAR positivity at the end of manufacturing when cultured with 100 IU IL-2/mL. 500 IU IL-2/mL n=5, 2 donors; 100 IU IL-2/mL uPAR-*TRAC*-CAR n=8, *TRAC*-Control n=6, 3 donors. (B) Summary of flow cytometry of %CAR and %mCherry expression with cultured conditions predicted to increase %CAR (100IU IL-2 and 0 μL CD3/CD28/CD2 activator after electroporation). uPAR-*TRAC*-CAR (blue, n=6), *TRAC*-Control (grey n=6) across three donors. (C) Mean Fluorescence intensity (MFI) of mCherry of both uPAR-*TRAC*-CAR and *TRAC*-Control T cells before and after directed fratricide with predicted culture conditions. uPAR-*TRAC*-CAR (blue, n=7), *TRAC*-Control (grey, n=7) across three donors. (D) Summary of flow cytometry immunophenotyping showing percent positive of T cell differentiation states: T_SCM_ CD45RA+/CD62L+ and T_CM_ CD45RO+/CD62L+ for uPAR-*TRAC*-CAR T cells with high antigen exposure (500IU IL-2 and 25µl CD3/CD28/CD2 activator after electroporation) and low antigen exposure (100IU IL-2 and 0µl CD3/CD28/CD2 activator after electroporation). High exposure n=5 across 2 donors, low exposure n=6 across 3 donors. Significance was determined by (A, B, C) two-way ANOVA, (D) ordinary one-way ANOVA; *p≤0.05; ***p≤0.001;****p≤0.0001. ANOVA, analysis of variance.

### Directed fratricide improves purity and potency after cryopreservation

Cryo-thawed CAR-T cells have demonstrated significantly elevated expression of mitochondrial dysfunction, apoptosis signaling, and cell cycle damage pathways^54,55^, and maintaining high cell yield and potency after cryopreservation is an essential feature of T cell therapies. Improvements in cryo-thaw would advance both allogeneic and autologous workflows, providing time - several days or even months - for quality release testing and coordination with clinical teams for infusion into the patient. To see if directed fratricide would improve cryo-thaw, we manufactured donor-matched uPAR-*TRAC*-CAR and muPAR- *TRAC*-CAR T cells without the directed fratricide abilities (see Figure 1H), and cryo-preserved cell products on day 9 of manufacturing. The cell products were then thawed after several weeks in liquid nitrogen storage and evaluated via flow cytometry (Figure 4A). After thawing, cells were immediately assayed for cell count and viability (Figure S6A-B). As expected, a modest decrease in viability and cell count was observed post-thaw due to cellular stress, resulting in early apoptosis. Upon further culture with IL-2 and activators supplemented media uPAR-*TRAC*-CAR T cells recovered robustly and maintained ≥99% CAR and TCR knockout two weeks post-cryopreservation. In contrast, the %CAR in the cryo-thawed muPAR-*TRAC*-CAR cells decreased over time in culture (Figure 4B; S6C).

**Figure 4.**
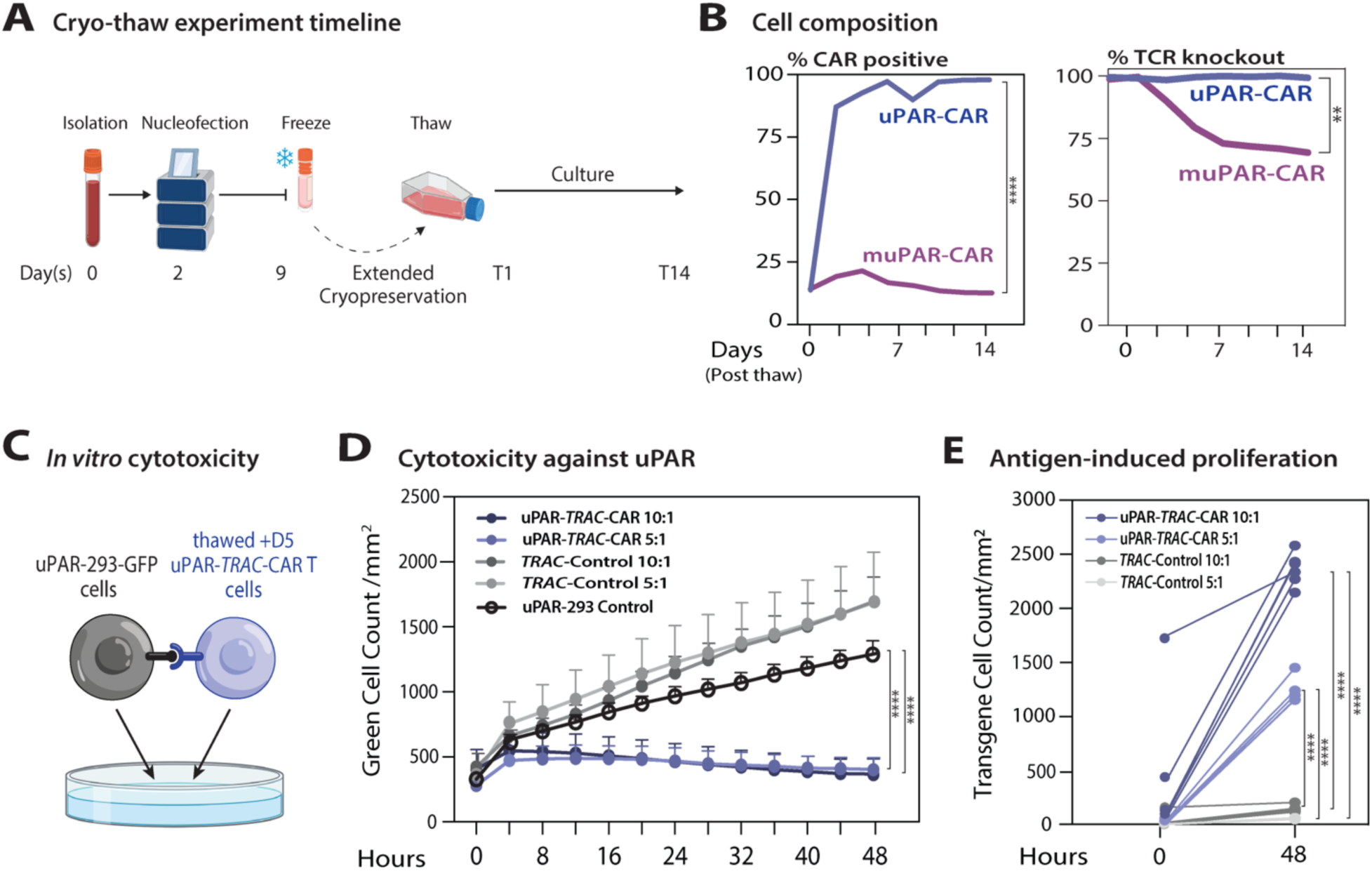
Cryo-thawed uPAR-*TRAC*-CAR T cells undergo directed fratricide and are potent. (A) *Ex vivo* manufacturing timeline for cryo-preservation and thawing of *TRAC*-CAR T cell products. (B) Flow cytometry summary data for transgene integration and TCR surface protein levels on the manufactured cell products on day 7 through 14 post-thaw for uPAR-*TRAC*-CAR (blue) and muPAR-*TRAC*-CAR (grey). (C) Schematic of uPAR-293 GFP co-culture assay with thawed day 5 uPAR-*TRAC*-CAR T cells. (D) IncuCyte® Live-Cell Analysis *in vitro* cytotoxicity of thawed uPAR-*TRAC*-CAR T cells against uPAR-293 GFP cells at effector:target ratio 10:1 and 5:1. The significant decrease in uPAR-293 GFP cells after T cells were added indicates high potency of uPAR-*TRAC*-CAR CAR T cells at both 10:1 and 5:1 effector: target ratios, uPAR-*TRAC*-CAR T cell 10:1 (dark blue) n=6, uPAR-*TRAC*-CAR T cells 5:1 (light blue) n=6, *TRAC*-Control 10:1 (dark grey) n=6 *TRAC*-Control 5:1 (light grey) n=6, uPAR-293 GFP (black) n=6. (E) IncuCyte® Live-Cell Analysis summary of Red Cell Count for mCherry positive cells at multiple effector:target ratios uPAR-*TRAC*-CAR blue (n=6), *TRAC*-Control (n=6). Significance was determined by ordinary one-way ANOVA; *p≤0.05; **p≤0.01; ***p≤0.001. ANOVA, analysis of variance.

To assess the potency of our thawed CAR T cell products, we measured cytotoxicity against uPAR-293 cells that had been genetically modified to express GFP (Figure 4C; S6D). We manufactured donor-matched uPAR-*TRAC*-CAR and *TRAC*-Control cells to act as a comparison to our fresh CAR T cell co-cultures. Our results showed significant cytotoxicity against the uPAR-293 cells for thawed uPAR-*TRAC*- CAR T cells compared to *TRAC*-Control cells (Figure 4D). Furthermore, we observed growth in the number of CAR-positive mCherry cells in the same experiments (Figure 4E). These results demonstrate that cryo-thawed uPAR-*TRAC*-CAR T cells retain the directed fratricide capabilities and can proliferate into potent cell products with high %CAR.

### Purifying an anti-GD2 CAR using directed fratricide

Our success in limiting antigen exposure and enhancing memory in uPAR-*TRAC*-CAR T cell populations compelled us to explore the potential of using this strategy to increase the purity of CAR T products directed against other clinically relevant antigens. We developed a dual targeting CAR (Dual-*TRAC*-CAR) that targets both uPAR and disialoganglioside (GD2), a tumor-associated surface antigen on many solid tumors^56,57^. The Dual-*TRAC*-CAR template contains a second-generation uPAR-*TRAC*-CAR co-expressed with a third-generation GD2-*TRAC*-CAR, with the costimulatory domains of CD28, OX40, and CD3σ (Figure 5A; S7A). In parallel as a control cell product, we also manufactured GD2-*TRAC*-CAR cells using the same virus-free methods. We achieved consistent genome editing with the dsDNA templates across three different donors and demonstrated up to 23% knock-in efficiency, with an average of 13% CAR+ and >90% total TCR- cells, as measured by flow cytometry (Figure 5B). We attribute the decrease in integration with the dual construct to the increase in the size of the donor template, which can limit overall knock-in efficiency^3,5^. Genomic integration of Dual-*TRAC*-CAR was further confirmed via PCR on genomic DNA from the cell product, with primers specific to the transgene (Figure S7D).

**Figure 5.**
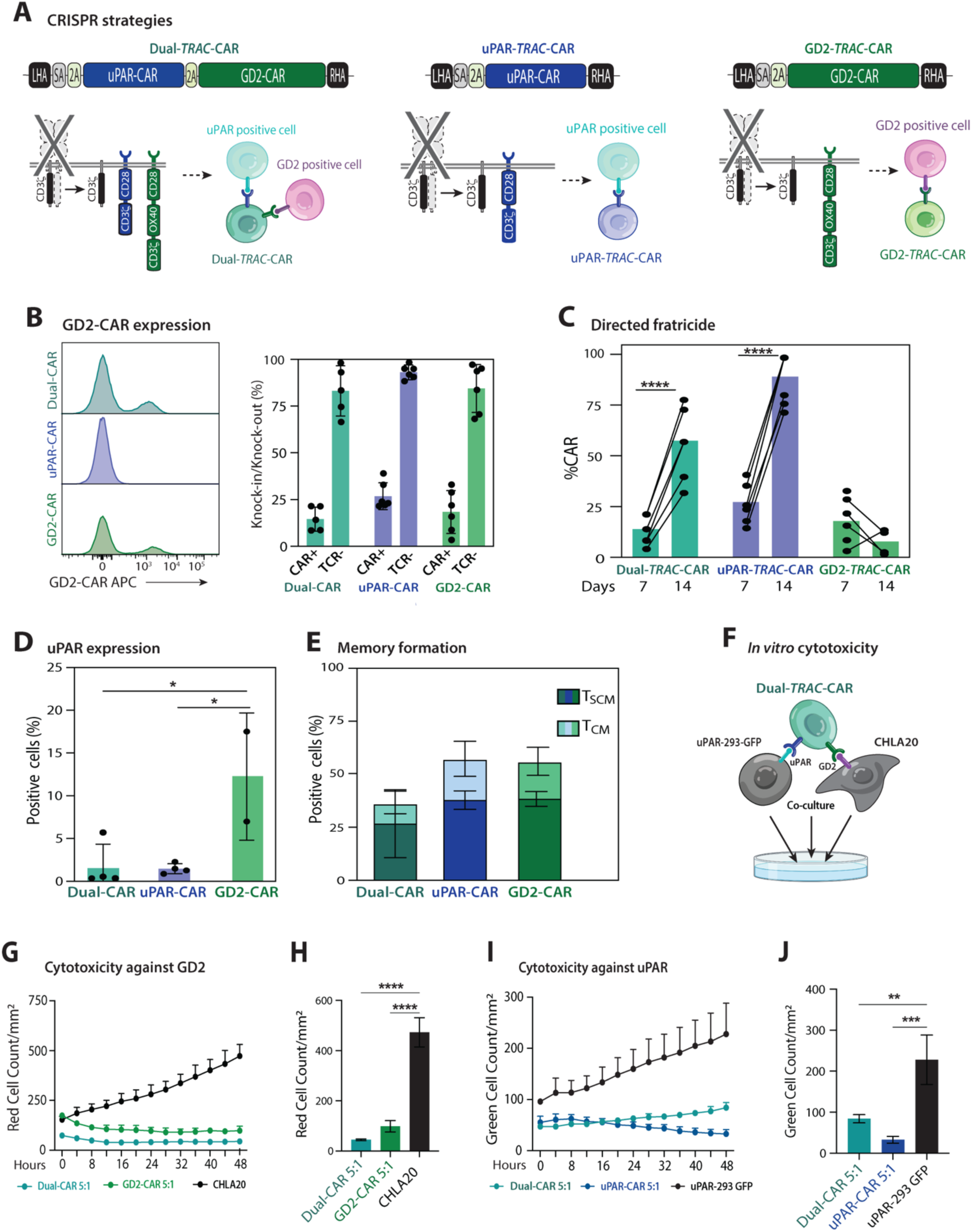
Purification of anti-GD2 CAR via directed fratricide. (A) Schematic of uPAR-*TRAC*-CAR, GD2-*TRAC*-CAR, and Dual-*TRAC*-CAR constructs targeting *TRAC*. uPAR-CAR (blue), GD2-CAR (green), and Dual-CAR (teal). (B) Histogram of GD2-CAR expression and summary of flow cytometry for knock-in and knock-out of the manufactured cell products on day 7 post-isolation. Dual-*TRAC*-CAR (teal, n=5), uPAR-*TRAC*-CAR (blue, n=6), GD2-*TRAC*-CAR (green, n=6), across two donors. (C) Flow cytometry for CAR in T cell cultures before and after a seven day directed fratricide period: day 7 and 14 post-isolation. Dual-*TRAC*-CAR (teal n=5), uPAR-*TRAC*-CAR (blue, n=6), GD2-*TRAC*-Control (green=6) across two donors. (D) Flow cytometry for uPAR expression after the directed fratricide period on day 14 post-isolation. Dual *TRAC*-CAR (teal n=4), uPAR-*TRAC*-CAR (blue, n=4), GD2-*TRAC*-Control (green=2) across one donor. (E) Flow cytometry immunophenotyping summary showing percent positive of T cell differentiate states: T_SCM_ CD45RA+/CD62L+ and T_CM_ CD45RO+/CD62L+ for both Dual-*TRAC*-CAR (teal, n=5), uPAR-*TRAC*-CAR (blue, n=5) and GD2-*TRAC*-CAR (green, n=5) on day 14 post-isolation across two donors. (F) Schematic of Dual-*TRAC*-CAR T cells with both uPAR-293 (GFP-positive) cells and CHLA20 (RFP-positive) cells. (G-H) IncuCyte® Live-Cell Analysis total Red Cell Count of Dual-*TRAC*-CAR T cells and GD2-*TRAC*-CAR against CHLA20s at 5:1 a effector:target ratio over 48 hours. Dual and GD2-*TRAC*-CAR T cells show significant cytotoxicity at 5:1 ratio (n=3). (I-J) IncuCyte® Live-Cell Analysis of total Green Cell Count of Dual *TRAC*-CAR T cells and uPAR-*TRAC*-CAR against uPAR-293 at a 5:1 effector:target ratio over 48 hours. Dual- and uPAR-*TRAC*-CAR T cells show cytotoxicity at 5:1 ratio (n=3). Significance was determined by (E, G, I) two-way ANOVA, (D, H, J) ordinary one-way ANOVA, (C) mixed ANOVA ;*p≤0.05; ***p≤0.001;****p≤0.0001. ANOVA, analysis of variance.

To test if our uPAR-CAR could increase %GD2-CAR in the Dual-*TRAC*-CAR system, we assayed the cells at day 7 and day 14 as described previously. We observed a significant increase in both Dual-*TRAC*-CAR and uPAR-*TRAC*-CAR T cell populations compared to the GD2-*TRAC*-CAR control (Figure 5C), up to 78% in %GD2-CAR without impacting overall growth relative to our controls (Figure S7B-C). On day 14, we additionally observed a significant decrease in the percentage of uPAR-positive cells in Dual-*TRAC*- CAR and uPAR-*TRAC*-CAR T cells compared to GD2-*TRAC*-CAR T cells, indicating that the directed fratricide was uPAR-antigen specific (Figure 5D).

At day 14, Dual-*TRAC*-CAR T cells displayed phenotypic profiles similar to uPAR-*TRAC*-CAR and GD2- *TRAC*-CAR cells (Figure 5E). Finally, we evaluated the cytotoxicity of our Dual-*TRAC*-CAR T cells against both uPAR-positive and GD2-positive cells through a co-culture assay. In this assay, GFP-positive uPAR-293 cells and RFP-positive, GD2-positive, CHLA20 neuroblastoma cells were seeded at a 1:1 ratio (Figure 5F). Live IncuCyte^TM^ imaging of the target cells showed significant cytotoxicity against GD2- positive cells at a 5:1 effector:target ratio for both Dual- and GD2-*TRAC*-CAR T cells (Figure 5G-H). Similarly, we observed cytotoxicity against uPAR-positive cells for both Dual and uPAR-*TRAC*-CAR T cells (Figure 5J-I). This data demonstrates that the Dual-*TRAC*-CAR cells were able to undergo directed fratricide to enrich for the desired potent cells and retain full GD2-target-specific cytotoxicity.

## Discussion

Here, we harness CRISPR genome editing and CAR T signaling to selectively target and eliminate uPAR- expressing cells. Our rapid, one-step non-viral manufacturing with autonomous directed fratricide presents a simplified biomanufacturing strategy, offering advantages such as precise genomic integration and pro-stem cell memory-like phenotypes. The specificity and cytotoxicity of these cells is consistent with other CAR T cell products^5,37,38^, including senolytic targeting CAR T cells produced using viral methods^27,28,32^.

In addition to its use as a standalone CAR, the uPAR-*TRAC*-CAR platform may be co-expressed alongside other CARs, such as CD19 or CD22, in a dual configuration. These multi-CAR strategies could enable directed fratricide of heterogeneous CAR T cell products, improving product consistency and scalability across diverse manufacturing workflows. Different CAR architectures, such as tandem ones^58^ and those with different co-stimulatory domains^59^, also might be needed for other antigens with differing antigen density from GD2 on the tumor cell surface. Moreover, the uPAR-CAR strategy could be adapted to other cell types, particularly γδ T cells^60,61^, cord blood derived T cells^62^, or natural killer (NK) cells^63,64^, which may not require additional MHC editing. This flexibility positions our directed fratricide strategy as potentially programmable into many cellular chassis.

The novel directed fratricide program within uPAR*-TRAC*-CAR T cell products and offers a versatile platform for generating scalable autologous cell therapies, where bead-based separation steps and viral vector manufacturing are unnecessary. Furthermore, allogeneic cell therapies could be advanced through additional genome editing to knockout MHC class I and class II molecules. A variety of genome editing strategies, including zinc-finger-based, TALE-based and CRISPR-based editors (nucleases, base editors, primer editors, and epigenetic editors), have been used to modify multiple genes to make hypoimmune cells, including strategies that knockout *B2M* and *CIITA* and overexpress CD47 "don’t eat me" signals, to produce cells that persist within animal models and patients for weeks to months. Combining the directed fratricide strategy with these approaches is likely to eliminate dysfunctional cells within 48-hours of uPAR-CAR expression as observed in our live cell imaging assays and within 7-days of cryo-thaw. With these capabilities, we anticipate significant gains in potency and efficacy *in vivo* after infusion of cryo-thawed uPAR-CAR and Dual-CAR cell products.

Overactivation of CAR T cell products can result in cytokine release syndrome or reduced efficacy^65,66^. In contrast, our uPAR-*TRAC*-CAR T cells maintain a resting memory phenotype following directed fratricide. To achieve more precision in controlling the cell types within the product, further control of both uPAR induction and suppression could be achieved. Refining culture supplements further with cytokines or small molecules further could modulate uPAR dynamics and downstream CAR levels underpinning directed fratricide. Such cell-intrinsic control of cell composition could help prevent premature activation prior to infusion and serve as a manufacturing tool to consistently generate products within the therapeutic window for patient dosing. muPAR-CAR cell products established through viral vector transduction could be dosed safely via systemic infusion into mice^27,28^.

Our evaluation of uPAR-*TRAC*-CAR T cell products was limited by small-scale benchtop manufacturing (<2.5 × 10⁷ cells per run). However, protocols for large-scale production using the other electroporation systems have been previously established and are compatible with our non-viral uPAR-CAR editing approach^5^. While we demonstrated efficient editing and purification with the Dual-*TRAC*-CAR system, further optimization is needed to enrich anti-GD2 CAR populations to ≥99% purity. Additionally, the decreased population of CD45RA^high^/CD62L^high^ cells in Dual-*TRAC*-CAR T cells is consistent with tonic signaling reported in prior studies^37^. In future work, nonlinear multivariate modeling of the Dual-CAR manufacturing process could be combined with other AI-methods^67–69^ to further provide in-line control of fratricide during manufacturing^70^, and guide the adaptation of culture conditions. Building upon the strategy described here, further programming and establishing tighter control over cell-cell interactions during manufacturing opens up a new design space to leverage fratricide for the productions of more potent cell therapy products.

## Materials and methods

### Cell lines

HDFa (Fibroblasts) were purchased from ATCC and maintained in DMEM high glucose (Gibco) supplemented with 10% FBS (Gibco) and 1% penicillin–streptomycin (Gibco). HMC3 (Microglia) were purchased from ATCC and maintained in EMEM (Gibco) supplemented with 10% FBS and 1% penicillin– streptomycin. 3T3 cells were maintained in DMEM and supplemented with 10% FBS and 1% penicillin-streptomycin. uPAR-293 and uPAR-293-GFP cells are HEK-293 stably transfected with the entire coding region of uPAR cDNA cloned in the SgFI-MluI site of the pCMV6-Empty vector^71^. Stably transfected cells were grown in DMEM supplemented with 10% FBS and 0.6-1.4 mg/mL Geneticin. Selection for transgene-positive cells was confirmed by fluorescence-activated cell sorting for human uPAR expression (ThermoFisher; clone VIM5, TF, cat: 17-3879-42) and/or GFP (>90%+). Cell lines were maintained in culture at 37°C in 5% CO_2_ and tested routinely to rule out mycoplasma contamination.

### Lipofection

To generate uPAR expressing HEK-293 cells, the ThermoFisher Lipofectamine 3000 reagent was used. Before transfection with the pCMV6 vector containing uPAR cDNA, HEK-293 cells were plated in a 6-well plate such that they were 70-90% confluent the next day. Twenty-four hours later, transfections were started by first diluting 7.5 μL of Lipofectamine 3000 in Opti-MEM (Gibco) to reach a total volume of 125 μL. While this solution was incubating at room temperature for 5 minutes, 2.5 μg uPAR-pCMV6 vector DNA was diluted in a solution containing 5 μL of P3000 reagent and Opti-MEM to reach a total volume of 125 μL. This 125 μL mixture containing the uPAR-pCMV6 vector was moved to the tubes containing 125 μL of diluted Lipofectamine 3000. This solution was incubated for 15 minutes at room temperature, and then the 250 μL was transferred to the HEK-293 cells in the 6-well plate. These cells were visualized over the next 4 days before undergoing selection with 700ug/mL geneticin.

### Isolation of primary T cells from healthy donors

This study was approved by the Institutional Review Board of the University of Wisconsin-Madison (#2018-0103), and informed consent was obtained from all donors. Peripheral blood was drawn from healthy donors into vacutainers containing heparin and transferred to sterile 50 mL conical tubes. Primary human T cells were isolated using RosetteSep Human T Cell Enrichment Cocktail (STEMCELL Technologies). T cells were counted using a Countess II FL Automated Cell Counter (ThermoFisher Scientific) with 0.4% Trypan Blue viability stain (ThermoFisher Scientific) at a 1:1 dilution. T cells were cultured at a final density of 2 million cells/mL in ImmunoCult-XF T cell Expansion Medium (STEMCELL) supplemented with 200 IU/mL IL-2 (Peprotech) and stimulated with 25µL/mL Human CD3/CD28/CD2 T cell Activator (STEMCELL) immediately after isolation, per the manufacturer’s instructions.

### Isolation of primary T cells from Leukocyte Reduction System (LRS) Cone

Human PBMCs were obtained from whole blood from anonymous donors from Versiti Blood Center (Milwaukee, Wisconsin USA) as an alternative to drawing from healthy donors. Briefly, PBMCs were flushed out of the cone with a 16G blunt needle and dilution medium (PBS + 2% FBS) into a sterile 50 mL conical tube. Primary T cells were isolated using RosetteSep Human T Cell Enrichment Cocktail (STEMCELL Technologies) and placed on a rocker for 20 minutes at room temperature. The solution was then diluted 1:1 with dilution medium and 30 mL was carefully layered on top of 15 mL of Lymphoprep (STEMCELL Technologies) solution in a SeptMate^TM^-50 (IVD) tube (STEMCELL Technologies). Tubes were centrifuged for 1200 g x 20 minutes. The top layer containing leukocytes was gently poured off into a new sterile 50 mL conical tube and spun at 300 g x 10 min and washed twice until the supernatant was clean. T cells were counted using a Countess II FL Automated Cell Counter (ThermoFisher Scientific) with 0.4% Trypan Blue viability stain (ThermoFisher Scientific) at a 1:1 dilution and resuspended as above.

### T cell culture

T cells were cultured in ImmunoCult-XF T cell Expansion Medium at a density of 1 million cells/mL after activation. After 24 hours post-electroporation, all *TRAC*-CAR T cells and *TRAC*-Control T cells were transferred without centrifugation to 1 mL of fresh culture medium with either 500 or 100 IU/mL IL-2. T cells were passaged, counted, and adjusted to 1 million cell per mL in fresh medium + IL-2 on days 5, 7, 9, 11, and 14 and onward after isolation. To cryopreserve cells, T cells were centrifuged after determining cell count on day 9, the media supernatant was removed and cells were resuspended in 500µl FBS. Next, the resuspended T cells were placed in cryovials and 500µL 20% DMSO+FBS was added. T cells were immediately placed in a Mr. Frosty^TM^ (ThermoFisher) and placed in the -80℃ overnight before being moved to liquid nitrogen for long term storage. To thaw cells, cryovials were placed in a 37℃ water bath for 1 minute and then diluted in 9 mL ImmunoCult-XF T cell Expansion Medium. T cells were then spun at 200 rcf for 3 minutes and counted using a Countess II FL Automated Cell Counter with 0.4% Trypan Blue viability stain at a 1:1 dilution. T cells were cultured at a final density of 2 million cells/mL in ImmunoCult-XF T cell Expansion Medium (STEMCELL) supplemented with 200 IU/mL IL-2 (Peprotech) and stimulated with 25 μL/mL Human CD3/CD28/CD2 T cell Activator (STEMCELL). For conditions requiring treatment with dasatinib (MedChemExpress), a 10mM stock was thawed and diluted in culture media to reach a concentration of 100 nM which was then administered at 1 μL per mL of culture media unless otherwise noted.

### Nanoplasmid templates

Plasmids were generated by Genscript by inserting our CAR constructs into a Nanoplasmid backbone. Nanoplasmid backbones include two components: a small (∼300 bp) R6K origin of replication and an RNA-OUT cassette (∼70 bp). To linearize CAR dsDNA HDR template, a restriction digest of the Nanoplasmid construct using SspI-HF (catalog no. R3132S, NEB) was performed. Following guidance from Tommasi *et al.*^5^, four restriction digest batch reaction were performed in 1.5-mL Eppendorf tubes (50 μL Nanoplasmid at 2 mg/μL, 125 μL CutSmart buffer, 25 μL SspI-HF enzyme, and 1,050 μL DNase-free water for 1,250 μL total). The digest was run at 37°C for 60 minutes and then heat inactivated at 65°C for 20 minutes according to the manufacturer’s instructions. To purify the digested Nanoplasmid product, the samples were pooled into 600 μL reactions for Solid Phase Reversible Immobilization (SPRI) cleanup (6X) using AMPure XP beads according to the manufacturer’s instructions (Beckman Coulter). Each of the 600 μL starting products was eluted into 30 µL of water. Bead incubation and separation times were increased to 5 minutes, and elution time was increased to 15 minutes at 37°C to improve overall yield. PCR products from round 1 cleanup were then subjected to an additional round of SPRI cleanup to ensure purity. Template concentration and purity were quantified using an IMPLEN NanoPhotemeter® N50. Concentrated template products were diluted in UltraPure H_2_O at a concentration of 2.5 µg/ μL according to Nanodrop measurements. For the Dual *TRAC*-CAR Nanoplasmid an additional Gencircle backbone and Cas-CLIPT^5^ cut site were added to the sequence allowing it to self-cleave and linearize during electroporation eliminating any further steps of purification once received from the manufacturer for subsequent electroporation.

### SpCas9 RNP preparation

RNPs were produced by complexing *TRAC* sgRNA (IDT) to SpCas9. In brief, sgRNA at a concentration of 50 µM was purchased from IDT and aliquoted in UltraPure water for each electroporation replicate. Recombinant sNLS-SpCas9-sNLS Cas9 (Aldevron, 10 mg/mL, total 0.8 μL) was added to the complexed gRNA at a 1.2:1 molar ratio and incubated for 15 minutes at 37°C to form an RNP.

### T cell nucleofection

Following guidance from the protocol developed by *Mueller et. al.*^37^, RNPs and HDR templates were electroporated 2 days after T cell isolation and stimulation. T cells were centrifuged for 3 minutes at 200g and counted using a Countess II FL Automated Cell Counter with 0.4% Trypan Blue viability stain (ThermoFisher). 1 million T cells were aliquoted and centrifuged for 10 min at 90 rcf. During cell spin, 2 μL of (2 µg/μL) HDR template (total 4 µg) per condition were aliquoted to PCR tubes, followed by RNPs and were incubated for at least 10 minutes. After the cell spin, supernatants were removed by vacuum, and cells were resuspended in 80 µL P3 buffer (Lonza), then transferred to PCR tubes containing RNPs and HDR templates, bringing the total volume per sample to 24 µl. Each sample was transferred directly to a 20 µL Nucleocuvette^TM^ Vessel. T cells were electroporated with a Lonza 4D Nucleofector with X Unit using pulse code EH115. Immediately after nucleofection, 80 µl of pre-warmed recovery medium with 1.5µM M3184, either 500 IU/mL IL-2 or 100 IU/mL IL-2, and either 0, 25, or 50 µL/mL ImmunoCult CD3/CD28/CD2 activator was added to each cuvette. Cuvettes were rested at 37°C in the cell culture incubator for 15 minutes. After 15 minutes, cells were moved to a round bottom 96 well plate with additional recovery media.

### Flow cytometry analysis

T cells were stained and analyzed every 7 days for CAR (mCherry or anti-GD2) and TCR expression as well as exhaustion/memory markers, using fresh or frozen cells. Briefly, T cells were stained for live/dead with GhostDye^TM^ Red780 (Cytek Biosciences), then blocked with Human Trustain FcX^TM^ (Biolegend), and then full stained in BD Brilliant Stain Buffer (BD Biosciences) with antibody mix. Flow cytometry was performed on an Attune NxT flow cytometer (ThermoFisher Scientific). GD2 CAR was detected using 1A7 anti-14G2a idiotype antibody (National Cancer Institute Biological Resources Branch) and conjugated to APC with the Lightning-Link APC Antibody Labeling kit (Novus Biologicals). The following fluorophore-conjugated antibodies were used for immunophenotyping (BL: Biolegend; TF; ThermoFisher): anti-human TCR ɑ/β BV421 (clone IP26, BL, cat: 306722), anti-human CD45RO FITC (clone UCHL1, BL, cat: 304204), anti-human CD45RA BV711 (clone HI100, BL, cat: 304138), anti-human CD62L BV650 (clone DREG-56, BL, cat: 304832), anti-human PD-1 FITC (clone A17188B, BL, cat: 621612), anti-human TIM-3 BV711 (clone F38-2E2, BL, cat: 345024), anti-human LAG-3 PE (clone 11C3C65, BL, cat: 369306), anti-human uPAR APC (clone VIM5, TF, cat: 17-3879-42). Downstream analyses of all spectral cytometry data were performed in FCS Express 7 Software.

### In-out PCR

Following guidance from *Mueller et. al.*^37^, genomic DNA was extracted from 100,000 cells per condition using DNA QuickExtract (Lucigen), and incubated at 65°C for 15 min, 68°C for 15 min, and 98°C for 10 min. Genomic integration of the CAR was confirmed by in-out PCR using a forward primer upstream of the *TRAC* left homology arm ATCTTGTGCGCATGTGAGGGGC, and a reverse primer binding within the CAR sequence GCAAGCCAGGACTCCACCAACC. PCR was performed according to the manufacturer’s instructions using Q5 Hot Start Polymerase (NEB) using the following program: 98°C (30 s), 35 cycles of 98°C (10 s), 67°C (20 s), 72°C (2 min), and a final extension at 72°C (2 min).

### Ionizing Radiation

Adherent Fibroblasts and microglia were irradiated 100 cm from the source to the cells using a Xstrahl RS225 irradiator then returned to a 37 °C, 5% CO_2_ incubator. The dose-rate was 2 Gy/min for the X-ray irradiator.

### SA-β-Gal

SA-β-gal staining was performed using CHEMICON Cellular Senescence Assay Kit (cat. KAA002 Millipore Sigma) at a pH 6.0 for human cells. Adherent cells plated in a 12 well plate and fixed with 500 μL Fixing solution (Millipore Sigma) and incubated at room temperature for 10 minutes, washed twice with 1X PBS and stained with freshly prepared 1X SA-β-gal Detection Solution (Millipore Sigma) at 37°C, without CO_2_ and protected from the light and left overnight. The SA-β-gal Detection Solution was removed and the cells were washed twice with 1X DPBS. Three high power fields were captured per well.

### IncuCyte *In Vitro* Assays

A total of 5,000 or 10,000 target cells (irradiated fibroblasts, irradiated microglia, uPAR-293, uPAR-293-GFP) were seeded in quadruplets per condition in a 96 well flat bottom plate. Forty-eight hours later, T cells were added to each well at various effector:target ratios. The IncuCyte S3 Live-Cell Analysis System (Sartorius, Catalog No 4647), stored at 37°C, 5% CO2. Images were taken every 4 hours for 48 hours. Green object count was used to calculate the number of uPAR-293 cells in each well with either cell tracker or GFP. Green object count was also used to calculate the number of irradiated cells, however, using Annexin V (Sartorius), an early apoptosis marker. Red object count was used to calculate the number of uPAR-*TRAC*-CAR T cells. Fluorescent images were analyzed with IncuCyte Base Analysis Software. In IncuCyte experiments with uPAR-*TRAC*-CAR only (Figure S1; 4), the percentage of CAR+ cells was not normalized between conditions, and *TRAC*-Control cells had a lower total percentage of CAR+ cells; in the second set of experiments with the Dual-*TRAC*-CAR, the percentage of CAR+ cells was used to calculate the effector:target (E:T) ratio for all assays.

### *In Vivo* Toxicity Assay

5xFAD mice gifted from the Ulland Lab were crossbred with C57BL/6NTac.Cg-*Rag2^tm1Fwa^ Il2rg^tm1Wjl^*mice (Taconic; cat: 4111-M). Male and female Rag2/Il2rg^-/-^5xFAD (Rag-5xFAD) mice (15-17 weeks old) were anesthetized during the procedure and fastened using a stereotactic instrument (KOPF Model 940). The hair was removed, then an incision was made at midline of the head. A 1-mm burr hole was drilled in the skull at proper position (1.0 mm lateral, 0.5 mm posterior from bregma). Through this hole, up to 500,000 muPAR-*TRAC*-CAR T cells were suspended in 5 μL PBS and injected into the lateral ventricle at 2.5 mm deep from the surface of skull over 5 min using 10 µL Microliter Syringe Model 85 RN, Small Removable Needle, 26s gauge, 2 in, point style 2 syringe (Hamilton). Mice were measured for weight loss, temperature using infrared thermography, and overall survival every day. At the end of the study mice were necropsied; brains and spleens were isolated and evaluated for tumor growth.

### Nonlinear Multivariate Analyses

Nonlinear regression was performed using Evolved Analytics DataModeler software to further investigate culture conditions and memory/exhaustion markers with increased uPAR-CAR expression. Linear pairwise correlations were evaluated using the CorrelationExplorer function. Rewarding simplicity and accuracy, linear and nonlinear algebraic models were simultaneously generated in independent evolutions using DataModeler’s SymbolicRegression function. The models were plotted as a function of fit (1 − *R^2^*) and complexity. Models with high accuracy (*R^2^*≥ 0.9) and low complexity ≤ 150 were further processed to ensure all top-level terms passed an ANOVA test of p<0.005, as well as an interval arithmetic test to ensure no singularities. The selected models were then analyzed using the VariablePresence and VariableCombinations functions to identify the dominant variables and variable combinations. Finally, the models involving the top variable combinations were aggregated into individual model ensembles using the CreateModelEnsemble function. The models in the ensemble agree at observed data points but diverge in extrapolated parameter space. The divergence provides a trust metric and defines a predictive model that best fits the imported dataset. Key statistical attributes of the ensembles were evaluated using the ModelSummaryTable function and prediction performance was assessed via the EnsemblePredictionPlot. The model ensembles were further evaluated using the ResponsePlotExplorer function to visualize the expression of uPAR-*TRAC*-CAR as a function of each individual variable and their interactive effects within the ensemble and the ResponseComparisonExplorer function was used to compare different variable combinations.

### Statistical analysis

Unless otherwise specified, all analyses were performed using GraphPad Prism (V.8.0.1). For optimization experiments with more than two groups, a one-way ANOVA was used, with the Prism recommended post-test. For editing and cytotoxicity experiments with multiple donors, a one-way or two-way ANOVA was used, followed by the Prism recommended post-test. Error bars represent mean (SD). ns, p ≥ 0.05; ∗p ≤ 0.05; ∗∗p ≤ 0.01; ∗∗∗p ≤ 0.001; ∗∗∗∗p ≤ 0.0001.

## Ethics Approval

All human studies were conducted using a University of Wisconsin-Madison IRB-approved protocol (2018-0103, approved 08/31/23). All animal experiments were approved by the University of Wisconsin-Madison Animal Care and Use Committee (ACUC) under protocol #M005915.

## Data availability

The datasets used and/or analyzed during the current study are available from the corresponding author on reasonable request.

## Acknowledgements

We acknowledge funding from the Wisconsin Research Forward Program (MS, KS), NIH/NCI R01 CA278051, the Grainger Institute for Engineering at UW-Madison (KS), and NIH R35 GM119644-01 (KS), and T32GM135119 (LS). We acknowledge the University of Wisconsin Carbone Cancer Center Flow Cytometry Laboratory, and 1S10RR025483-01 (BD FACS Aria II BSL-2 Cell Sorter), for use of its facilities and services. We also acknowledge the Biomedical Research Model Services core, for the use of its facilities and services We also thank members of the Saha lab for helpful input on experimental design and manuscript preparation. The contents of this article do not necessarily reflect the views or policies of the Department of Health and Human Services, nor does mention of trade names, commercial products, or organizations imply endorsement by the US Government.

## Author contributions

LS designed all sequences. LS designed, performed, and analyzed knock-out experiments and sequencing studies. LS and DG collected whole blood, isolated T cells, and established the cell expansion and gene editing protocol. LS, DG, CS, RTr, MN generated dsDNA template. LS, DG, AU, BK, NM designed, performed, and analyzed gene editing optimization studies and co-culture experiments. TK performed analytical modeling. LS and RTa performed all animal experiments. LS, DG, and KS wrote the manuscript with input from all authors. KS supervised the research. All authors read and approved the final manuscript.

---

**Figure S1.**
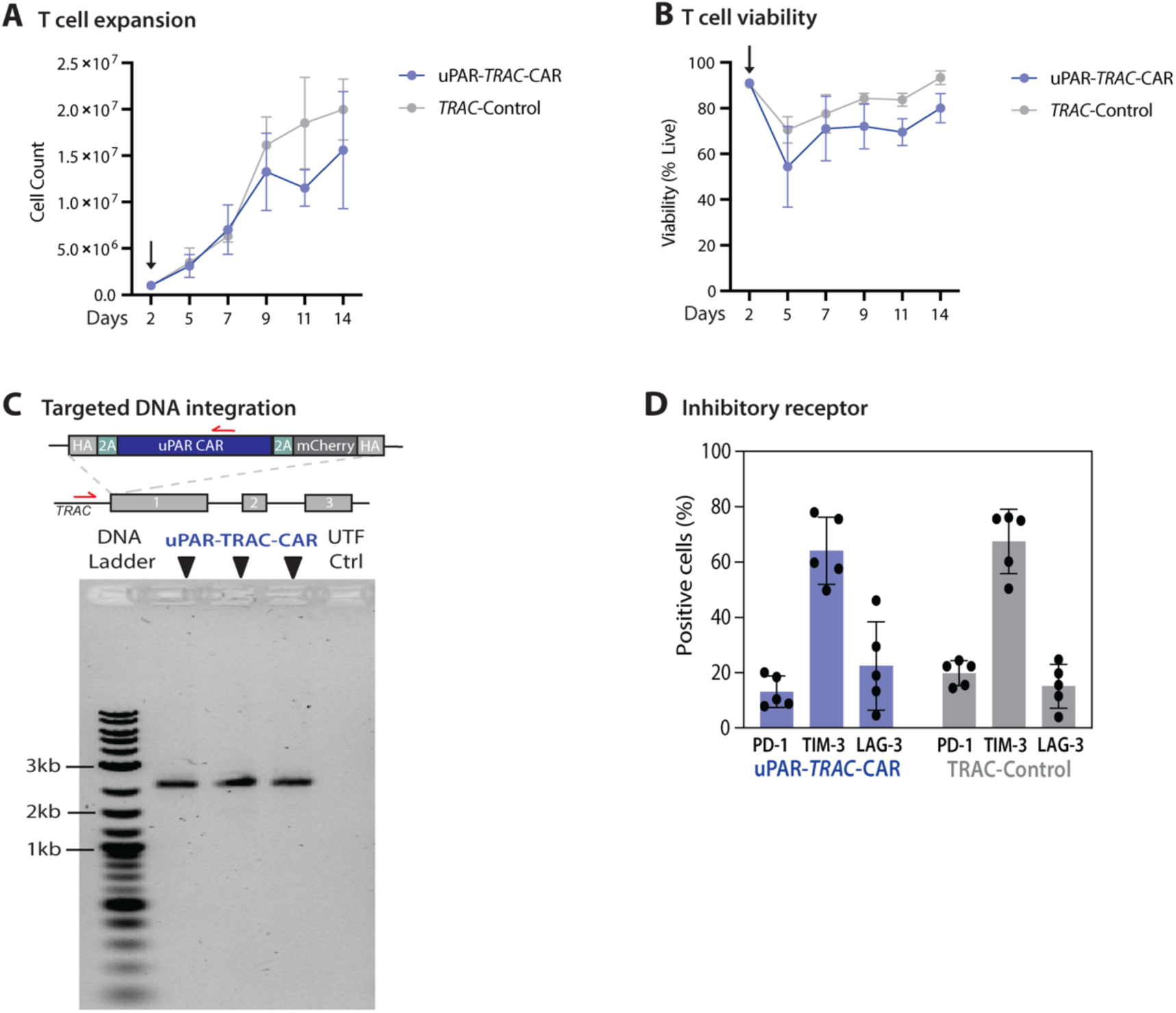
Characterization of uPAR-*TRAC*-CAR T cells. (A-B) Cell expansion and viability of uPAR-*TRAC*-CAR T cells throughout the 14 day *ex vivo* manufacturing process; uPAR-*TRAC*-CAR (blue) n=5, *TRAC-*Control (grey) n=5 across three donors. (C) Integration PCR, 2.6kb band indicates proper on-target genomic integration of the CAR transgene in uPAR-*TRAC*-CAR CAR cells and absent in UTF, untransfected donor-matched T cells. Primer locations (red arrows) are upstream of the left homology arm and within the CD28 sequence of the CAR to confirm on-target CAR transgene integration. This band was absent in the untransfected T cell donor matched controls. (D) Summary of flow cytometry for exhaustion markers (inhibitory receptors) PD-1, TIM-3, and LAG-3 for uPAR-*TRAC*-CAR (blue, n=5) and *TRAC*-Control (grey, n=5) on day 14 post isolation across three donors.

**Figure S2.**
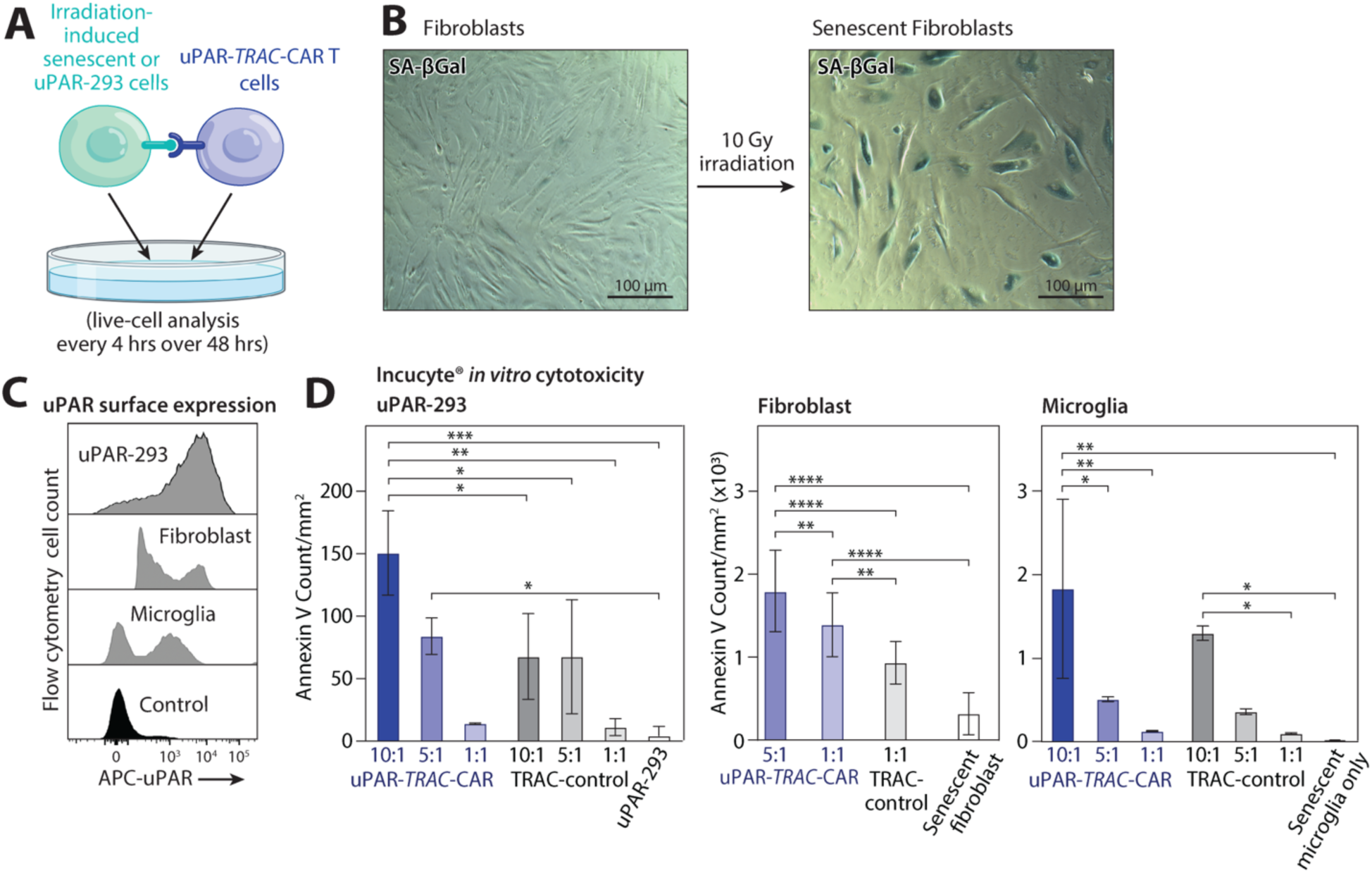
Cytotoxicity of uPAR-*TRAC*-CAR T cells against senescent induced and uPAR-positive cells. (A) Schematic of senescent induction of Human Dermal Fibroblasts (HDFa), human iPSC-microglia (iPSC-MG) with 10Gy irradiation and then validated for uPAR protein expression. (B) Cytochemical staining of SA-ß-gal in normal proliferating HDFas (Control) or after 10 Gy irradiation. (C) Flow cytometry histograms for uPAR surface expression on a human embryonic kidney (HEK) control cell line, uPAR overexpressing HEK line created by liposome-based transfection (uPAR-293), and on human dermal fibroblasts and microglia cells after 10Gy irradiation to induce senescence. (D) IncuCyte® Live-Cell Analysis cytotoxicity of uPAR-*TRAC*-CAR against senescent induced fibroblasts, microglia, and uPAR-293 cells at various effector:target ratios. The significant increase in the number of apoptotic cells after T cells at the end of coculture (48 hours) indicates high potency of uPAR-*TRAC*-CAR T cells. uPAR-*TRAC*-CAR (blue) n=6; *TRAC*-Control (grey) n=6; across two donors. Significance was determined by ordinary one-way ANOVA *p≤0.05; **p≤0.01; ***p≤0.001; ****p≤0.0001. ANOVA, analysis of variance.

**Figure S3.**
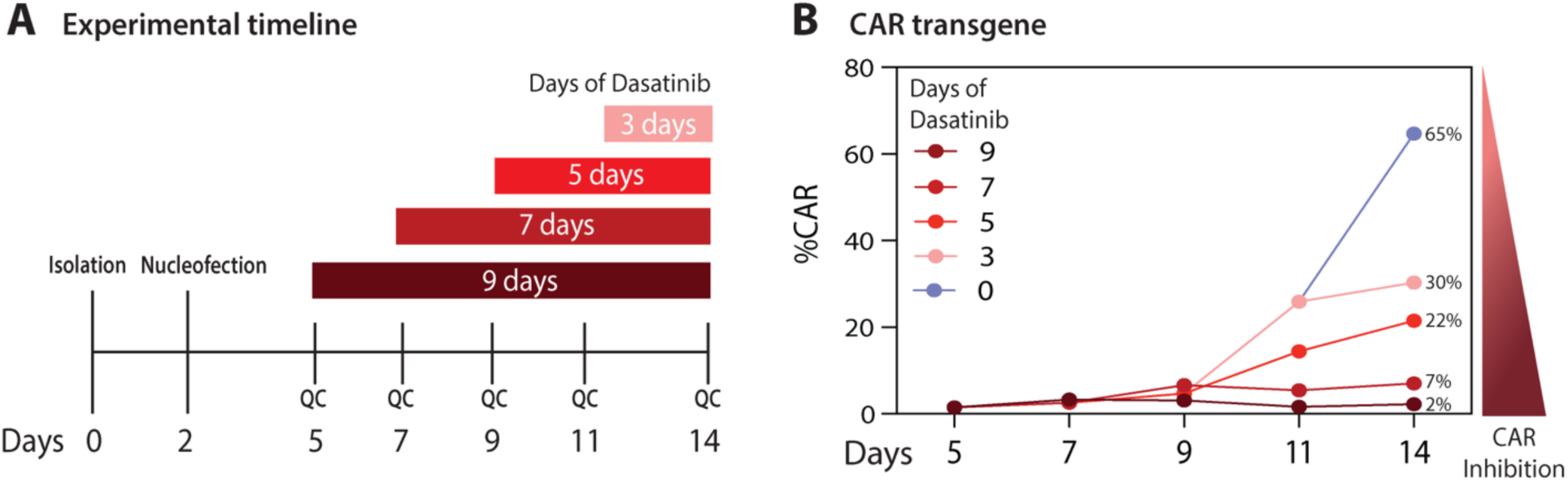
Dasatinib suppresses uPAR-*TRAC*-CAR expression. (A) *Ex vivo* manufacturing timeline of culture uPAR-*TRAC*-CAR T cells with dasatinib. Duration of dasatinib treatment is shown starting on day 5 post T cell isolation (maroon), day 7 (dark red), day 9 (red), day 11 (pink); “QC” = quality control flow cytometry. (B) Flow cytometry results for uPAR-*TRAC*-CAR %CAR expression throughout dasatinib treatment. Earlier treatment after nucleofection with dasatinib suppressed directed fratricide resulting in decreased %CAR expression (Control; uPAR-*TRAC*-CAR without dasatinib treatment (blue)).

**Figure S4.**
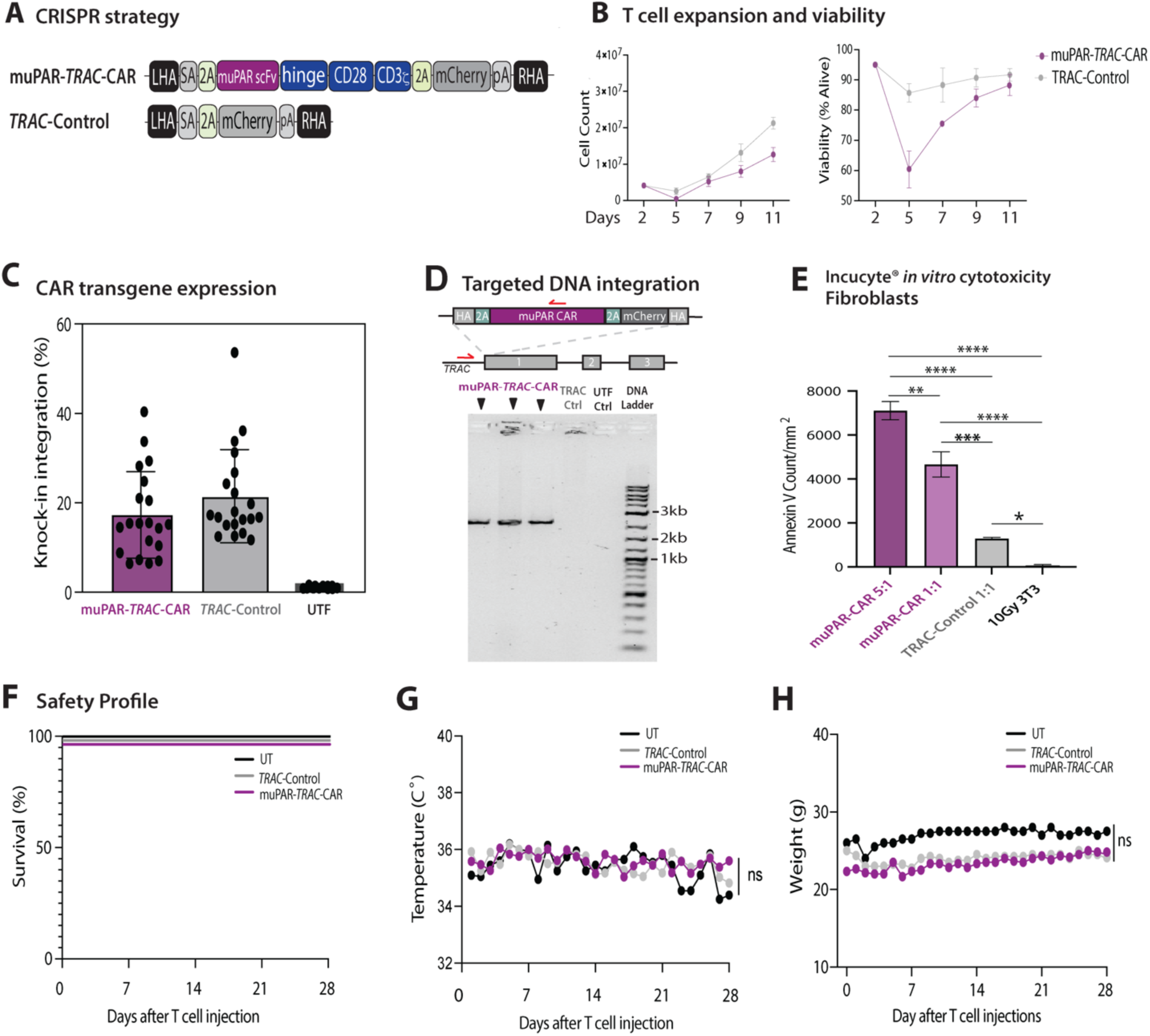
Generation and safety profiling of muPAR-*TRAC*-CAR T cells. (A) Schematic of muPAR-*TRAC*-CAR construct targeting using the first encoding exon of the human *TRAC* locus, SA: splice acceptor (grey), 2A: self-cleaving peptide (light green), uPAR scFv: single chain variable fragment targeting murine uPAR (purple), hinge domain, human costimulatory domains: CD28 and CD3z (blue), 2A: self-cleaving peptide (light green), mCherry: fluorescent tag protein (grey), pA: rabbit ß-globin polyA terminator (grey). (B) Cell expansion and viability of muPAR-*TRAC*-CAR T cells throughout the manufacturing; muPAR-*TRAC*-CAR (purple) n=3, *TRAC*-Control (grey) n=3 across two donors. (C) Flow cytometry plots for transgene expression muPAR-*TRAC*-CAR T cell products. Y-axis shows knock-in integration on day 7 post-isolation. muPAR-*TRAC*-CAR (purple, n=20), *TRAC*-Control (grey, n=20), Untransfected (UTF) (black, n=20) across 6 donors (D) Integration PCR, 2.6kb band indicates proper on-target genomic integration of the CAR transgene in muPAR-*TRAC*-CAR T cells. Primer locations (red arrows) are upstream of the left homology arm and within the CD28 sequence of the CAR to confirm on-target CAR transgene integration. *TRAC*-Control, and UTF, untransfected donor-matched T cells are controls. (E) IncuCyte® cytotoxicity of muPAR-*TRAC*-CAR T cells against senescent induced murine fibroblasts (3T3) at effector:target ratio 5:1 and 1:1. The significant increase in the number of apoptotic cells after T cells were added indicates high potency of muPAR-*TRAC*-CAR T cells at both 5:1 and 1:1 effector: target ratios, muPAR-*TRAC*-CAR T cell 5:1 (purple) n=3, muPAR-*TRAC*-CAR T cells 1:1 (light purple) n=3, *TRAC*-Control 1:1 (grey), senescent 3T3s (black) n=3. (F-H) *In vivo* safety profile of immunodeficient mice infused with muPAR-*TRAC*-CAR T cells. (F) Kaplan–Meier curve showing event-free survival of mice after treatment with either muPAR-*TRAC*-CAR T cells, *TRAC*-Control T cells, or Untreated PBS (UT) (muPAR-*TRAC*-CAR, n = 6; *TRAC*-Control, n = 5 mice; UT, n=3). (G-H) Weight (G) and temperature (H) of mice every 24 h after T cell infusion throughout the duration of the 28 day study (muPAR-*TRAC*-CAR, n = 6; *TRAC*-Control, n = 5; UT, n = 3 mice). Significance was determined by ordinary one-way ANOVA **p≤0.01; ****p≤0.0001.

**Figure S5.**
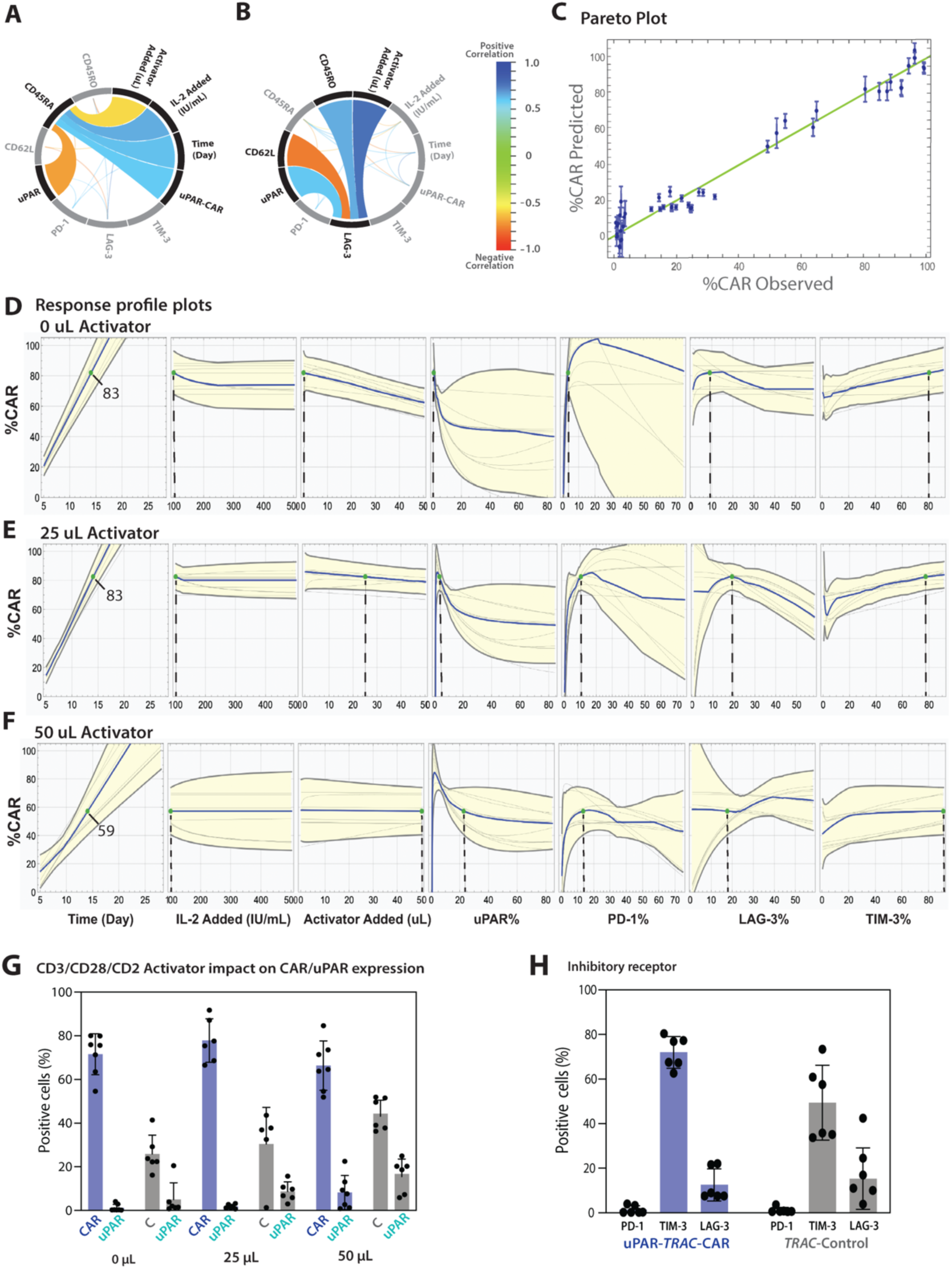
Nonlinear regression analysis shows a relationship between culture conditions, %CAR, and exhaustion. (A-B) Variable correlation plots for (A) CD45RA and (B) LAG-3. (C) Model ensemble prediction plot demonstrating high predictive power of the model ensemble for the %CAR observed. The error bars identify the 2σ spread of the ensemble predictions for each data point (D) Response profile plots of the model ensemble for uPAR-CAR with 0 μL activator for time, IL-2, activator, uPAR, and exhaustion. (E) Response profile plots of the model ensemble for uPAR-CAR with 25 μL activator for time, IL-2, activator, uPAR, and exhaustion. (F) Response profile plots of the model ensemble for uPAR-CAR with 50 μL activator for time, IL-2, activator, uPAR and exhaustion. The gray lines represent individual models, while the blue line represents the median prediction of the ensemble. The yellow shows the 2σ spread of the ensemble predictions. (G) Flow cytometry of %CAR/mCherry and %uPAR expression for uPAR-*TRAC*-CAR (blue) and *TRAC*-Control (grey) on day 14 manufacturing after culture with 0µl (0x), 25µl (1x), or 50µl (2x) Immunocult Human CD3/CD28/CD2 T cell activator and 100 IU IL-2/mL. There was a notable increase in mCherry and uPAR expression for *TRAC*-Control with increased activator; no difference in mCherry expression from day 7 to 14 for *TRAC*-Control observed (data not shown). n=8, across 3 donors. (H) Flow cytometry of exhaustion markers (inhibitory receptors) PD-1, TIM-3, and LAG-3 for uPAR-*TRAC*-CAR (blue, n=6) and *TRAC*-Control (grey, n=6) on day 14 post isolation across 3 donors.

**Figure S6.**
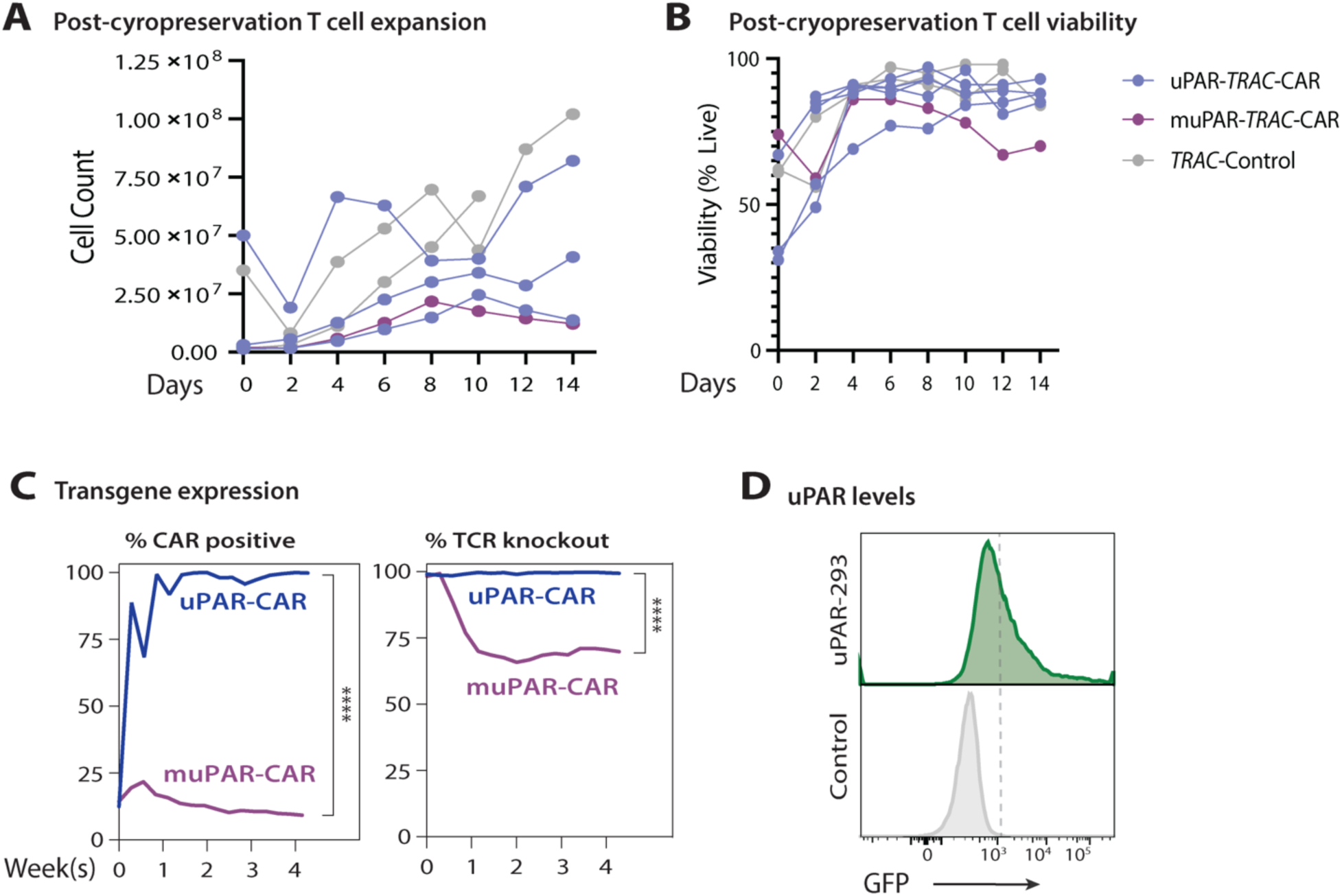
Cryo-thawed uPAR-*TRAC*-CAR T cell undergo cell expansion and long term directed fratricide. (A-B) Cell expansion and viability of uPAR-, muPAR-*TRAC*-CAR T cells, and *TRAC*-Control T cells after cryo-preservation and -thaw of *TRAC*-CAR T cell products; uPAR-*TRAC*-CAR (blue) n=5, muPAR-*TRAC*-CAR (purple) n=1, *TRAC*-Control (grey) n=3 across two donors. (C) Flow cytometry summary data for transgene integration and TCR surface protein levels on the manufactured cell products on day 7, 14, 21, and 28 post-thaw. (D) Flow cytometry histogram for GFP expression of uPAR-293 after lipofection with Lonza pmaxGFP™ vector GFP and Control untreated uPAR-293 cells. Significance was determined by Welch’s t test **p≤0.01; ****p≤0.0001. ANOVA, analysis of variance.

**Figure S7.**
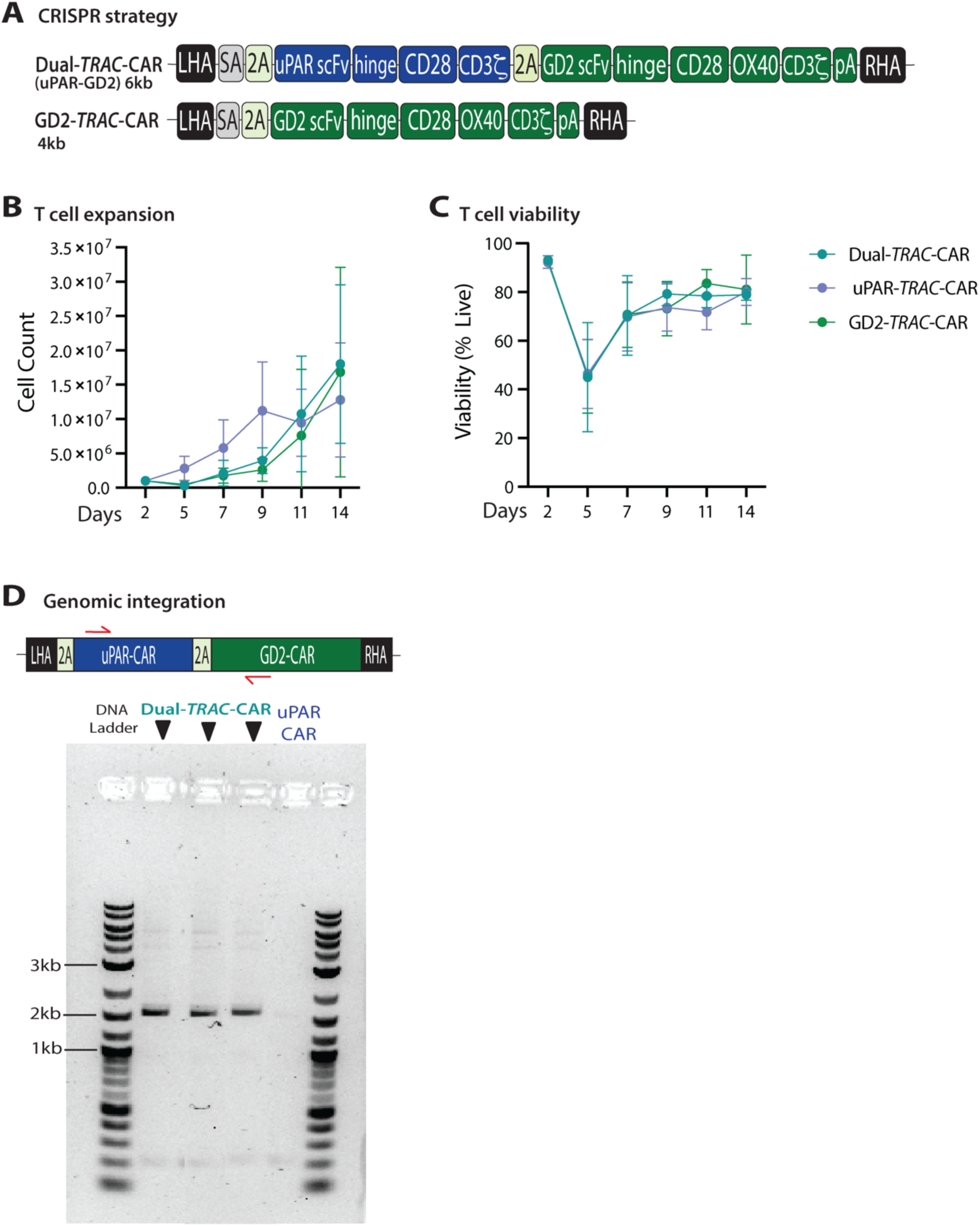
Characterization of Dual-*TRAC*-CAR T cells. (A) Schematic of Dual-*TRAC*-CAR and GD2-*TRAC*-CAR construct targeting using the first encoding exon of the human *TRAC* locus, SA: splice acceptor (light green), T2A: self-cleaving peptide (green), uPAR scFv: single chain variable fragment targeting human uPAR (blue), uPAR-CAR hinge domain, uPAR-CAR costimulatory domains: CD28 and CD3**ζ** (blue), P2A: self-cleaving peptide, GD2 scFV: single chain variable fragment targeting human GD2 (green); GD2 CAR hinge domain, GD2-CAR costimulatory domains: CD28, OX40, CD3**ζ** (green), pA: rabbit ß-globin polyA terminator (green). (B-C) Cell expansion and viability of Dual-, uPAR-, and GD2-*TRAC*-CAR T cells T cells throughout the 14 day *ex vivo* manufacturing process; Dual-*TRAC*-CAR (teal) n=6, uPAR-*TRAC*-CAR (blue) n=6, GD2-*TRAC*-CAR (green) n=6 across three donors. (D) Dual-CAR primer design (red arrows) and PCR showing integration of the Dual-CAR transgene in Dual-*TRAC*-CAR CAR T cells as indicated by the band at 2.1kb across three donors. This band is absent in the uPAR-*TRAC*-CAR T cell control.

